# Methylation-Guided Stratification of Colorectal Cancer Reveals Immune Subtypes with Distinct Clinical Outcomes

**DOI:** 10.64898/2026.02.03.703457

**Authors:** Eiman I. Ahmed, Raghvendra Mall, Christophe M. Raynaud, Heba Saadeh, Shimaa Sherif, Rania Alanany, Nady El Hajj, Davide Bedognetti, Jessica Roelands, Wouter Hendrickx

## Abstract

**Background:** Aberrant DNA methylation is a hallmark of colorectal cancer (CRC). Yet, how DNA methylation is linked to transcriptional states, immune programs, and tissue resident microbiome within the same tumors has not been systematically analyzed.

**Methods:** We profiled genome-wide DNA methylation (Illumina MethylationEPIC) in 182 colon tumors and 76 adjacent normals from AC-ICAM, and integrated with matched transcriptomes, whole exome, microbiome, and clinical data. Tumor-specific methylation, promoter methylation–expression links, microbiome associations, and survival were analyzed and validated in TCGA-COAD.

**Results:** Tumor and normal tissues exhibited distinct DNA methylation patterns, reflecting widespread epigenetic alterations in cancer. Pathway analysis identified two major tumor pathways regulated by DNA methylation. The first involved extracellular signaling and adhesion genes, with higher methylation linked to increased proliferation and lower immune infiltration. Similarly, higher tumor methylation in nitric oxide signaling was associated with reduced adaptive immune activity and interestingly, influenced immune-related survival. These findings were also validated in the TCGA-COAD cohort. An inverse methylation–expression pattern implicated modifications of TCR signaling in naïve CD8⁺, and interferon-α/β signaling which were hypermethylated and hypomethylated in tumors compared to normal, respectively. Combining methylation and microbiome revealed connections between *Akkermansia muciniphila* and TGF-β and *Prevotella nigrescens* with MAPK signaling pathways. Finally, a methylation-based model using 43 promoters CpGs successfully identified patients with different survival outcomes, underscoring the clinical relevance of these epigenetic alterations in colon cancer.

**Conclusion:** DNA methylation shapes the molecular and immune landscape of colon cancer, altering signaling pathways and immune programs, interacting with the microbiome, and impacting patients’ survival.

## Introduction

Colorectal cancer (CRC) is among the most commonly diagnosed malignancies worldwide and a leading cause of cancer mortality. Its development reflects a layered interplay between genetic and epigenetic remodeling that together reprogram epithelial identity and tumor–host interactions (Gu et al., 2019; Janssens et al., 2023). Within this continuum, aberrant DNA methylation stands out as a defining hallmark of CRC: it emerges early, accumulates through progression, and intersects with pathways that govern proliferation, invasion, apoptosis, and immune surveillance (Baharudin et al., 2022).

Epigenetic architecture underlies well-recognized molecular strata in CRC. The CpG island methylator phenotype (CIMP) captures a broad tendency toward promoter hypermethylation, while microsatellite instability (MSI) reflects mismatch repair (MMR) deficiency that, in sporadic disease, frequently arises from promoter methylation of MMR genes. These features are not independent: CIMP-high tumors are often MSI-high and align with the immunogenic Consensus Molecular Subtype 1 (CMS1), highlighting the tight link between methylation patterns, genomic instability, and immune contexture in CRC (Joo et al., 2023; A. H. Li, 2024; J. Liu et al., 2019). Collectively, these insights underscore DNA methylation as an active driver of transcriptional regulation, antigen presentation, cytokine signaling, and clinically relevant tumor phenotypes, rather than simply a passive marker of transformation (Kumar et al., 2025).

Concurrently, host–microbiome interactions shape, and are shaped by, the colonic methylome. Microbial metabolites and inflammation can induce locus-specific methylation changes, while methylation-driven transcriptional programs remodel the epithelial niche and immune milieu, potentially selecting for distinct bacterial communities (Gutierrez-Angulo et al., 2023). Despite this reciprocal influence, integrated analyses that connect methylation states to immune features, microbiome composition, and patient outcome within the same cohort remain limited.

The AC-ICAM cohort provides a deeply annotated, publicly accessible multi-omics resource in CRC spanning exome, transcriptome, immune profiling, microbiome sequencing, and clinical annotations (Roelands et al., 2023). Here, we extend AC-ICAM with high-density, genome-wide DNA methylation profiling (Illumina MethylationEPIC array) in primary tumors and matched normal tissues. This study aims to systematically and comprehensively characterize DNA methylation in CRC by integrating it with the other omics layers from the AC-ICAM, to better understand tumor–immune–microbiome interactions, define methylation subtypes, and identify prognostic markers.

## Materials and methods

### Study cohort

Samples used in this study are part of the AC-ICAM cohort. Detailed description of patients’ selection and samples processing can be found in our previously published manuscript (Roelands et al., 2023). Here we used Infinium MethylationEPIC array to sequence 84 adjacent-normal and 194 colon tumor samples, in which 74 are paired normal-tumor samples.

### Infinium MethylationEPIC sequencing

Genome-wide DNA methylation analysis was performed using the Illumina Infinium MethylationEPIC BeadChip v1.0 (Illumina, San Diego, CA), which interrogates over 850,000 CpG sites at single-nucleotide resolution. Genomic DNA per sample was bisulfite-converted using the EZ DNA Methylation Kit (Zymo Research, Irvine, CA) according to the manufacturer’s instructions. Converted DNA was then amplified, enzymatically fragmented, and hybridized to the BeadChips following the Illumina Infinium HD Methylation protocol. Arrays underwent single-base extension and fluorescent labeling and were scanned using the Illumina iScan System. Methylated and unmethylated signal intensity data were generated as idat. files. The QC of the assay control probes was performed on the Illumina GenomeStudio software. All the downstream analyses were conducted using the hg19/GRCh37 human genome assembly.

### Methylation data processing

The raw idat files were converted to raw intensity data and stored in RGChannel object minfi R package (v. 1.46.0). Subsequent processing was done using mainly minfi R package following three steps: samples quality control (QC), normalization and probes QC. Samples QC was done based on the detection *P*-value which was calculated using ‘detectionP’ function. Samples with p-value less than 0.01 were excluded. Moreover, samples with median methylation intensity less than 2 x standard deviations (SD) were excluded. Samples with sex discrepancies between reported sex from clinical data and predicted sex from ‘getSex’ function were excluded. Data normalization was then performed using ‘preprocessFunnorm’ function with background and dye bias correction with noob and quantile extension. Probes QC was done by excluding cross reactive probes, probes on SNPs location (Chen et al., 2013, Pidsley et al., 2016), and probes located on sex chromosomes. In addition, ‘filter_impute_MMatrix’ function (Bergstedt et al., 2022) was implemented to exclude low quality probes. The M-matrix was converted to Beta (β) value matrix using the following equation:

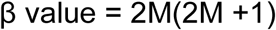

The final β matrix contained 258 samples (76 adjacent normal and 182 tumors) and 705,084 CpGs. Probes annotations were obtained from Illumina Infinium MethylationEPIC manifest file from IlluminaHumanMethylationEPICmanifest package (v.0.3.0). For CpG-gene mapping we used ‘UCSC_RefGene_Name’ column in the manifest file by extracting the first gene name of this column. For TCGA-COAD, raw idat files were downloaded from GDC using ’GDCquery‘ function from TCGAbiolinks R package (v. 2.28.3). The idat files were processed following the same pipeline used in AC-ICAM cohort.

### Dimension reduction

Dimension reduction based on methylation profile was performed using Uniform Manifold Approximation and Projection (UMAP). β value matrix was passed to ‘umap’ function from umap R package (v.0.2.10.0) using default settings. UMAP 1 and UMAP 2 coordinates were selected for plotting scatter plot using ggplot2 R package (v.3.3.5) to visualize differences in methylation patterns.

### Differentially methylated positions (DMPs)

DMPs analysis was conducted to compare two groups of interest using the ‘dmpFinder’ function from the minfi R package (v.1.46.0). The analysis was performed with default settings to identify DMPs using β matrix. The difference in β values (delta-beta (Δβ)) between the two groups was determined by computing the mean β value for each CpG site within each group and then subtracting the mean from one group from the other. Minimum Δβ cutoff of 0.3 was selected to compare the methylation difference and increased to 0.4 to define the most differently methylated CpGs between tumor and normal for pathway analysis. Genes with both hypomethylated and hypermethylated DMPs were filtered out. For the comparison between hyper- and hypo-methylated genes between tumor and normal, we computed the mean for all CpGs (FDR < 0.01, Δβ >= |0.4|) mapped to each gene and plotted the top 15 hypomethylated and top 15 hypermethylated genes in a heatmap to compare between tumor and normal. DMPs were plotted in a heatmap to visualize their methylation pattern using Bioconductor R package ComplexHeatmap (v. 2.16.0).

### Consensus clustering

To identify methylation subtypes within tumor samples, we performed consensus clustering on β value matrix using ConsensusClusterPlus (v.1.64.0) R package. Consensus clustering was performed for k value of 6 with 5000 resampling and agglomerative hierarchical clustering with ward criterion (Ward.D2) inner and complete outer linkage. We selected the clusters of two groups as we intended to cluster the groups into hypo- and hyper-methylated. The cluster with the higher β-mean for CpGs of interest was considered the hypermethylated cluster, while the group with lower β-mean was considered the hypomethylated group. For determining CIMP phenotype, we performed consensus clustering based on previously published CpGs by Joo et al (Joo et al., 2023) which consist of the following seven CpGs mapped to five genes: CACNA1G: cg18337803, cg20467136, cg23614229, cg11262815; RUNX3: cg06377278, cg27095256; SOCS1: cg06220235; NEUROG1: cg04620091; and IGF2: cg16977706. However, two CpGs (cg18337803, cg20467136) were included in SNPs locations, therefore they have been filtered out during the QC process, hence the clustering was performed without them. To determine CIMP subtypes (k = 3), the cluster with the highest promoter CpGs β-mean was labeled as CIMP high (CIMP-H), and the cluster with the lowest CpGs mean labeled as CIMP low (CIMP-L). Samples with intermediate CpGs β-mean were classified as CIMP medium (CIMP-M).

### RNA sequencing and processing

Normal and tumor samples were processed and normalized as described in detail in our previously published manuscript (Roelands et al., 2023). Normalized and log2 transformed gene expression matrix was used for all downstream analysis.

### Differentially expressed genes (DEGs)

DEGs analysis was performed using limma (v.3.56.2) Bioconductor R package with Benjamini-Hochberg (B-H) FDR. In each comparison, genes with rows sum equal to zero or no gene expression across samples were removed. Genes with chosen cutoff were used for downstream analysis as specified in each analysis.

### Pathway enrichment analysis

Lists of genes mapped to DMPs or list of DEGs were uploaded to consensus path DB (CPDB) website or Ingenuity Pathways Analysis (IPA) to get the list of enriched pathways. For CPDB, output data was then downloaded as an ORA file and loaded to R studio for analysis. Pathway clustering was then done using func2vis (v.1.0-3) R package (Orecchioni et al., 2022). For IPA, data associated with the pathways was exported in form of an excel file. Pathways that are not related to CRC were removed. Bar plots were generated using ggplot2 (v.3.3.5) package.

### Single samples gene set enrichment analysis (ssGSEA)

Single-sample gene set enrichment analysis (ssGSEA) was performed to calculate enrichment scores (ESs) using R package GSVA (v.1.38.2). Gene sets that correspond to particular tumor-associated pathways or immune-related signatures were curated from multiple sources as previously described in detail (Roelands et al., 2020) and (Roelands et al., 2023) respectively. A list of genes for leukocyte extravasation signaling was obtained from the molecular signatures database (MSigDB). ES for pathways derived from CPDB analysis were calculated using the genes mapped to the corresponding pathway from CPDB. To visualize the differences between groups of interest, ESs were displayed in boxplot using ggplot2 or aggregate median heatmap using ComplexHeatmap (v. 2.16.0).

### Gene Ontology

Gene Ontology (GO) enrichment analysis was performed to identify biological processes associated with the gene list of interest. First, the human gene annotation database org.Hs.eg.db from org.Hs.eg.db R package (v. 3.21.0) was used to map gene symbols to entrez gene identifiers. The conversion was performed using the ‘bitr’ function from the clusterProfiler package (v. 4.16.0). GO enrichment analysis was then carried out using the ‘enrichGO’ function from the clusterProfiler package, specifying org.Hs.eg.db as the annotation database. The parameters were set as follows: pvalueCutoff = 1, qvalueCutoff = 1, and minGSSize = 1, allowing an inclusive exploration of enriched terms. To reduce redundancy among significantly enriched GO terms, the results were simplified using the ‘simplify’ function from the clusterProfiler package. Data visualization was performed using barplot from enrichplot R package (v. 1.28.4).

### Survival analysis

Overall survival (OS) data was used from the clinical data to perform survival analysis and to generate Kaplan-Meier (KM) curves using ‘ggsurvplot’ function from survminer (v. 0.4.9) R package. Patients with less than one day of follow up were excluded and survival data were censored after a follow-up duration of 20 years. Hazard ratio (HR), *P*-value, and confidence intervals (CIs) were calculated using ‘coxph’ function from survival (v. 3.8-3) R package.

### Methylation risk-score model development

For model development and training, M-value matrix of 899 promoter CpGs with intermediate CpG-gene expression negative correlation (R >= -0.5) from AC-ICAM dataset was used. Since we aimed to use the TCGA-COAD as a validation cohort, we selected from the initial list of CpGs (n=899), a 700 promoter CpGs overlapping between the two cohorts for model development and validation.

We first normalized the M-value matrix by converting each CpG column into a *z*-score using the mean and SD and treating the normalized M-value matrix as the training set. We then built a multivariate elastic-net OS Cox regression model using the *glmnet* R package (v.4.1.9) on the training set. The optimal hyper parameters (λ) for the best model were identified through five-fold cross-validation via a grid-search technique using the ‘cv.glmnet’ function. We used the concordance index as a performance metric. The parameters for which the mean cross-validation concordance index was the highest were selected as optimal hyper parameters. Next, the final model was built using the optimal λ value on the complete training set. To calculate risk scores in the training dataset (methylation risk-scores), we passed the training set and best model to the ‘predict’ function from *glmnet* package. A total of 43 features (CpGs) were present in the best model with nonzero coefficients; we refer to these features as the ‘methylation classifier’, which comprise the final survival model. A positive or negative coefficient of the M-value for each CpG of the methylation classifier can be binarized into ‘high-risk’ and ‘low-risk’ groups using the cutoff threshold of 0 and attributed to the strength of association with survival. A higher positive coefficient means high hazards or risk of death, whereas a negative coefficient corresponds to lower risk of death.

#### Methylation risk-score model validation

We validated the final model on the TCGA-COAD cohort. The TCGA-COAD consists of 181 samples. We processed the M-values for each CpG by converting the M-value matrix into *z*-scores using the mean and SD derived from the training set of AC-ICAM. This normalized M-value matrix was passed to the predict function along with the best model to estimate corresponding risk scores. The risk score (methylation score) of any tested sample is only dependent on the M-value of the list of CpGs with nonzero coefficients in the methylation classifier (the risk score for each sample is not dependent on one of the other samples). Finally, the methylation scores are binarized using the cutoff threshold 0 to categorize the test sample into ‘high-risk’ (>0) and ‘low-risk’ (<0) groups as performed in the training set. Therefore, no cutoff optimization occurred in the validation phase.

#### Methylation risk-score model performance assessment

We tested the concordance index (CI) in the training set using the final methylation model; (1) in the training set using the cross-validation of the best methylation model (ten permutations, 80% training and 20% validation partition) achieving a CI of 0.579 ± 0.020; and (2) the test set cohort of the TCGA-COAD, where it achieved a CI of 0.601.

### Statistical analysis

The statistical significance for the comparison of CpGs β-mean value, genes expression, enrichment scores, differences in genus and species abundance between groups of interests was calculated using unpaired Wilcoxon test using R function ‘stat_compare_means’ from ggpubr R package (v.0.6.0). Statistical significance was accepted for p < 0.05 and p < 0.01 or as specified in each analysis. Spearman correlation was performed using ‘cor.test’ function from R stats package. For CpG-species correlation, centered log-ratio (CLR) transformation was applied to the β matrix before performing the correlation using ‘clr’ function from compositions R package (v.2.0-9).

## Results

### AC-ICAM methylation profile

In this study, we expanded our publicly available AC-ICAM dataset by appending methylation data from EPIC Infinium array profiling. A total of 182 tumors and 76 neighboring healthy colon tissue methylomes were integrated with existing clinical and molecular data (**Figure 1A**).

**Figure 1:**
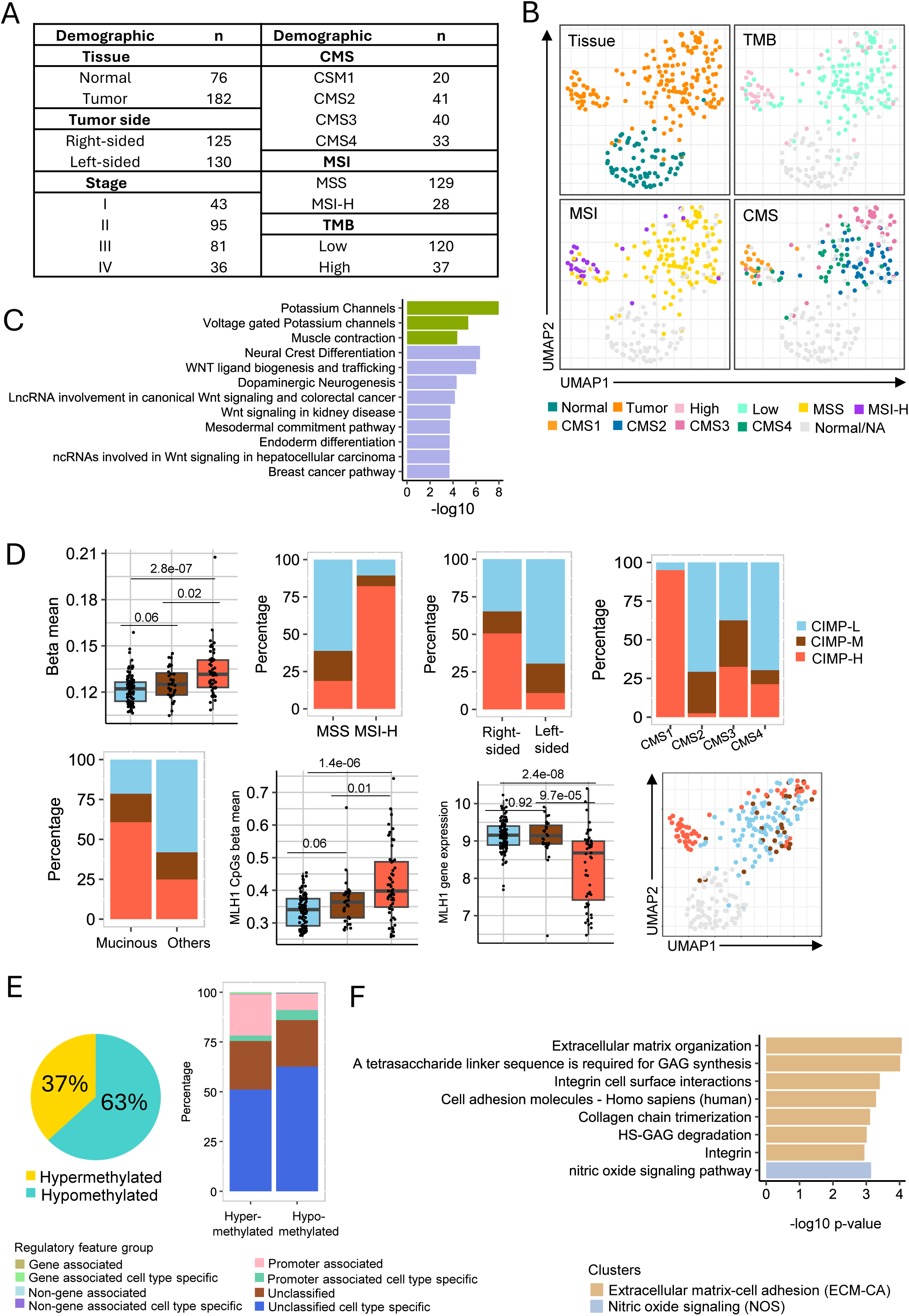
Overview of AC-ICAM methylation profile. **A.** Summary of AC-ICAM cohort for samples included in this study. **B.** UMAP plots based on methylation beta-value. Samples annotated with tissue, TMB, MSI status, and CMS. Grey dots TMB, MSI and CMS plots represent normal samples or missing data from tumor samples. **C.** Pathways of genes mapped to DMP (FDR < 0.01, Δβ >= |0.3|) between tumor-subgroup and other tumor samples. The plot shows the enrichment of pathways divided into two sets, each represented by a separate color. **D**. CIMP analysis. The top left boxplot shows the average beta-value of promoter CpGs in CIMP subtypes. CIMP-H has the highest average beta value, while CIMP-L has the lowest. Stack-bar charts show the frequency of CIMP subtypes in MSI groups, colon side, CMS subtypes and mucinous subtypes, respectively. CIMP-H has hypermethylated CpGs of MLH1 genes; a marker of CIMP subtypes. On the other hand, CIMP-L has hypomethylated MLH1. As a result, MLH1 expression was low in CIMP-H compared to CIMP-L. The UMAP plot annotated with CIMP shows that the tumor-subgroup has high enrichment of CIMP-H phenotype. Represented numbers in boxplots resemble the *P* value from Wilcoxon test. **E.** The pie chart demonstrates the differentially methylated positions (DMPs) between tumor and normal in which 63% of DMPs are hypomethylated in tumor, while 37% are hypermethylated when compared to normal (DMP FDR < 0.01, Δβ >= |0.3|). The stack-bar chart shows hyper/hypomethylated CpGs annotated with feature group. CpGs with no annotation in the manifest file were excluded. The hypermethylated CpGs have higher frequency of promoter CpGs. **F.** Pathway enrichment analysis using Consensus Path DB database (CPDB) for genes (n = 207) mapped to DMPs (n = 439, FDR < 0.01, Δβ >= |0.4|) between tumor and normal. The barplot shows two different sets of pathways, each represented by different colors namely, extracellular signaling-cell adhesion (ECM-CA) and nitric oxide signaling (NOS).

To gain a high-level perspective of the global differences in methylation profile between samples, we performed dimension reduction analysis using UMAP. Major differences in methylation profile were observed between tumor and normal tissues, which formed separate clusters in UMAP-embedding **(Figure 1B)**. Interestingly, we noticed a tumor subgroup enriched in high tumor mutation burden (TMB), high microsatellite instability (MSI-H), and Consensus Molecular Subtype 1 (CMS1) samples **(Figure 1B)**. This demonstrates the interconnection of methylation profiles with these molecular classifications. Further pathway enrichment analysis of genes (n = 485) mapped to differential methylated probes (DMPs) (*n* = 960, FDR < 0.01, Δβ-value >= 0.3) between the tumor subgroup and other tumor samples revealed enrichment of WNT signaling pathways (**Figure 1C**).

Next, to explore CpG island methylator phenotype (CIMP) we performed consensus clustering for 7 CpGs (Joo et al., 2023) from previously described genes that are known to be associated with CIMP (Weisenberger et al., 2006). Consensus clustering was able to successfully classify the tumor samples into three CIMP subgroups CIMP-H, CIMP-M and CIMP-L (**Supplementary figure 1A**). Tumors with CIMP-H demonstrated, as expected, extensive hypermethylation across promoter CpGs when compared to CIMP-L (*P* = 2.8e-07) (top, left boxplot **Figure 1D**). Consistent with the literature (Joo et al., 2023; M. Li et al., 2023; J. Liu et al., 2019), the CIMP-H subgroup is enriched in MSI-H (*P* = 2.2e-10), right-sided colon (*P* = 3.2e-08), CMS1 (*P* = 6.8e-12) and mucinous samples (*P* = 0.0003). Moreover, CIMP-H is associated with *MLH1* hypermethylation when compared to CIMP-L (*P* = 1.4e-06), resulting in a lower *MLH1* gene expression (*P* = 2.4e-08). Similar results for association between CIMP, MSI, CMS and MLH1 were obtained in TCGA-COAD (**Supplementary figure 1 B-D**). We also observed a clear enrichment of CIMP-H in the identified subgroup of tumors that clustered separately on the UMAP (**Figure 1D**).

Next, to systematically identify the aberrant modifications in methylation status between tumor and normal samples across the whole genome, we performed DMPs analysis. A total of 158,862 DMPs (FDR < 0.01 with at least 0.3 in Δβ-value) were identified. DMPs with higher Δβ-value in tumor compared to normal were considered hypermethylated, whereas DMPs with lower Δβ-value were considered hypomethylated. Based on this, we noticed that 63% of the DMPs are hypomethylated and 37% are hypermethylated in tumor compared to normal (pie chart **Figure 1E**). Of note, a higher proportion of promoter CpGs (in pink and green) were present in the hypermethylated DMPs, indicating potential epigenetic gene silencing in tumor compared to normal tissue (stacked bar chart **Figure 1E**). When assessing methylation alterations between tumor and normal samples at the gene level, we identified 207 genes mapped to 439 DMPs (FDR < 0.01, Δβ-value >= 0.4). Among these, 52 genes exhibited hypomethylation, while 155 genes were hypermethylated in tumor samples based on the mean β-value of CpGs mapped to each gene (**Supplementary figure 1E**). To further understand the biological significance of the differences in methylation profiles between tumor and normal tissues, we performed pathway enrichment analysis on genes (n = 207) mapped to DMPs between tumor and normal samples (n = 439, FDR < 0.01, Δβ-value >= 0.4). Two different sets of curated pathways were significantly enriched in tumor samples (FDR < 0.05) (**Figure 1F**) (full list of pathways is available in supplementary table 3). The first set was represented by *e*xtracellular matrix organization, the most enriched pathway (-log10 p-value = 4.06, *P* = 8.65e-05), and included related processes such as integrin signaling and glycosaminoglycan (GAG) metabolism. Collectively, we refer to this set as the extracellular matrix and cell adhesion (ECM-CA). The second set reflects only “nitric oxide signaling” (-log10 p-value = 3.15, *P* = 7.16e-04), therefore we labeled this as nitric oxide signaling (NOS). To gain deeper insight into the biological significance of these findings, we focused on these two enriched sets of pathways by investigating their associated CpG methylation patterns in tumor versus normal tissues, and within tumor.

### Methylation Changes in Extracellular Matrix and Cell Adhesion Genes are linked to Tumor Proliferation and Immune Suppression

Given the significant enrichment of ECM-CA pathways in tumor-specific differentially methylated genes (**Figure 1F**), we examined the CpGs associated with these pathways to assess their methylation patterns and biological relevance in tumors. Consensus clustering of tumor samples based on the 27 CpGs associated with ECM-CA pathways stratified the samples into two distinct groups. ECM-CA high methylation (ECM-CA-HM) exhibits a higher mean β value for these CpGs (*P* < 2.2e-16) compared to ECM-CA low methylation (ECM-CA-LM). When comparing to normal samples, ECM-CA-LM displayed an intermediate methylation pattern, positioned between normal and ECM-CA-HM **(Figure 2A)**. A similar trend is also observed in TCGA-COAD (**Supplementary figure 2A**). The methylation pattern of the 27 CpGs reveals that only three CpGs have lower methylation in tumor samples compared to normal, while the remaining CpGs exhibit higher methylation. Notably, CpGs mapped to the same gene tend to cluster together. Additionally, differences in CpGs methylation pattern were observed within tumor samples between the ECM-CA-LM and ECM-CA-HM clusters, with the ECM-CA-LM cluster displaying a methylation pattern more closely resembling that of normal samples. Moreover, a higher enrichment of CIMP-H is observed within ECM-CA-HM (**Figure 2B**). An inverse correlation between methylation and gene expression has been observed for several genes such as *ITGA4, PXDN, VCAN, COL4A1* and *COL23A1* suggesting that epigenetic alterations contribute to the dysregulation of ECM-CA pathways in tumors (**Figure 2C, supplementary figure 2B**). Modifications in extracellular matrix components have previously been associated with tumor proliferation (Karlsson & Nyström, 2022; Z.-L. Li et al., 2020; Yuan et al., 2023), prompting us to investigate the enrichment of proliferation-related pathways. Our analysis revealed that the ECM-CA-HM cluster exhibits a higher mean proliferation enrichment score (ES) compared to ECM-CA-LM (*P* = 0.018) and normal samples (*P* < 2.2e-16). Notably, we observed the same trend in the TCGA-COAD cohort (*P* = 9e-05) (**Figure 2D**). To further explore the differences between ECM-CA clusters, we performed differentially expressed genes (DEGs) analysis between ECM-CA-HM and ECM-CA-LM. The list of significant genes (FDR < 0.01, log2FC ≥ |0.5|) was used for pathway enrichment analysis via Ingenuity Pathway Analysis (IPA). Among the top 20 most significant pathways, we observed a notable enrichment of immune-related pathways, which were downregulated in ECM-CA-HM compared to ECM-CA-LM. The z-score further confirmed the lower activity of these pathways in ECM-CA-HM. Notably, the most significantly enriched pathway was leukocyte extravasation signaling (*P* = 1.0e-15) (**Figure 2E**). To quantify this difference, we calculated an ES for this pathway, which highlights a marked reduction in ECM-CA-HM compared to ECM-CA-LM (*P* = 5.3e-11) and normal samples (*P* < 2.22e-16). A similar trend was observed in the TCGA-COAD cohort, where ECM-CA-HM exhibited significantly lower leukocyte extravasation signaling ES than ECM-CA-LM (*P* = 0.00017) (**Figure 2F**). While the ECM-CA clusters reflect meaningful biological variation, they were not associated with differences in overall survival (**Supplementary figure 2C**). Overall, these findings suggest that elevated ECM-CA methylation may reduce expression of key extracellular matrix and adhesion genes, potentially altering the ECM landscape in a way that limits leukocyte extravasation and immune cell infiltration. Collectively, tumors with higher ECM-CA methylation are characterized by increased proliferation and reduced immune pathway activity, indicating a potential epigenetic mechanism of immune evasion in a subset of colon cancers.

**Figure 2:**
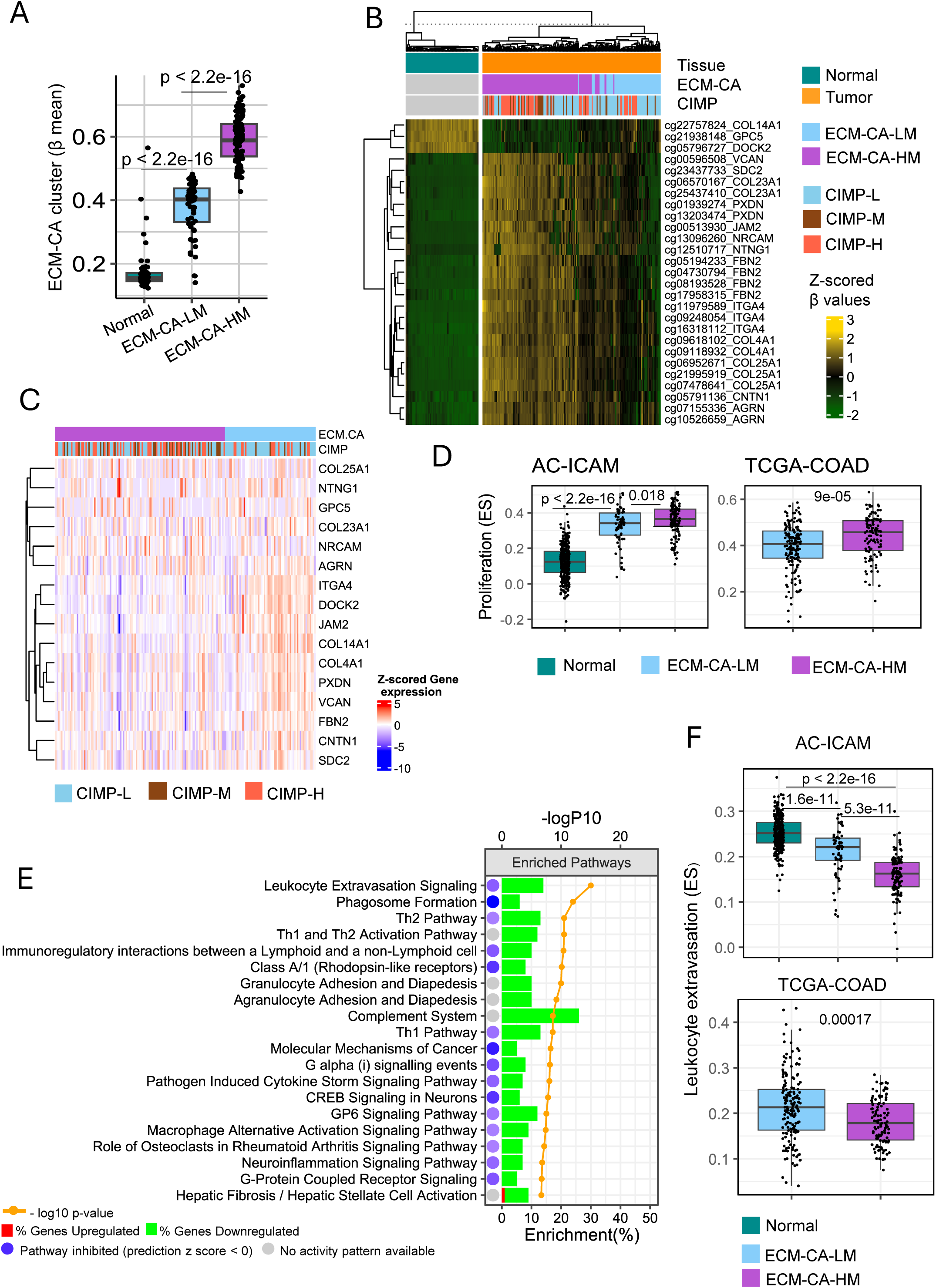
ECM-CA CpG high methylation defines a proliferative and immune-cold colon cancer subgroup. **A.** Boxplot shows the average β for 27 CpGs mapped to genes associated with ECM-CA DMPs from figure 1F for normal, ECM-CA-HM and ECM-CA-LM clusters. Plotted *P*-value from Wilcoxon test. **B.** Heatmap of the 27 DMPs from panel A using β-values. The heatmap is annotated by tissue, CIMP, ECM-CA LM/HM clusters based on consensus clustering method for the 27 DMPs. Dendrogram column clustering for tumor and normal samples were performed separately for each group. **C.** Heatmap of expression for genes (n = 16) mapped to the 27 CpGs from panel B. **D.** Boxplots comparing proliferation ES between normal, ECM-CA-LM and ECM-CA-HM in AC-ICAM (left plot) and TCGA-COAD (right plot). *P*-value represented from Wilcoxon test. **E.** Top 20 enriched pathways (based on *P*-value) for differentially expressed genes (DEG) (n = 520, FDR <= 0.01, log2FC >= |0.5|) between ECM-CA-HM and ECM-CA- LM clusters. Analysis was performed using IPA database. Histograms represent the proportion (%) of DEGs upregulated (red) or downregulated (green) in ECM-CA-HM vs. ECM-CA-LM. The circles represent the pathway activation status. Blue circle indicates the pathway is inhibited with a negative z-score, while a gray circle indicates that the pathway activity is unknown. The connected orange dots indicate the -log10 p-value for each pathway. **F.** Boxplots of AC-ICAM (top plot) and TCGA-COAD (bottom plot) showing the ES of leukocyte extravasation signaling; the top significant pathway from panel E. Statistical analysis based on Wilcoxon test when comparing between groups in boxplots.

### Nitric Oxide Methylation Define Immune Phenotypes and Stratify Survival Outcomes

Similarly, enrichment analysis identified nitric oxide signaling (NOS) as a significantly overrepresented pathway among genes associated with DMPs (**Figure 1F**). Tumor samples were stratified into two clusters using consensus clustering on eight CpGs mapped to the NOS pathway. These were defined as NOS high methylation (NOS-HM) and NOS low methylation (NOS-LM), with NOS-HM showing higher mean β values in AC-ICAM (*P* < 2.2e-16) (**Figure 3A**). Similar clusters were observed in TCGA-COAD (**Supplementary figure 2D**). The methylation patterns of these CpGs indicate that they all have higher methylation in tumors compared to normal samples, suggesting potential epigenetic gene silencing in tumor tissues (**Figure 3B**). A significant inverse correlation (R = -0.39, *P* = 1.2e-10) between NOS β mean and NOS ES based on all genes mapped to the pathway was observed (**Supplementary figure 2E**), supporting downregulation of these genes as consequence of gene methylation. Normal samples have significantly higher NOS ES compared to both NOS-LM and NOS-HM groups, while the difference between NOS-LM and NOS-HM is trending toward significance (*P* = 0.077) (**Figure 3C**). To gain insight into the biological differences between the NOS-HM and NOS-LM, we performed pathway enrichment analysis on DEGs (n = 940, FDR < 0.01, log2FC <= |0.5|). We observed that the downregulation of DEGs associated with several immune pathways such adaptive immune system, chemokines, TCR and BCR signaling pathways in NOS-HM compared to NOS-LM (top 10 pathways from each **Figure 3D, supplementary figure 3**). Given the immune-related findings and the role of NOS in shaping the tumor microenvironment, we integrated NOS ES to capture the functional activity of the pathway with the Immunologic Constant of Rejection (ICR) to assess their combined prognostic value. The ICR score, a signature derived from the mean expression of 20 immune-related genes, reflects the overall immune context of the tumor microenvironment (Roelands et al., 2020). We divided the patients based on NOS ES median into NOS ES high and NOS ES low and similarly for ICR score. While no significant difference in survival was observed between the NOS ES groups (top KM curve, **Figure 3E**), patients with ICR high demonstrated, as expected, significantly better survival outcomes compared to the ICR low group (*P* = 0.009) (bottom KM curve, **Figure 3E**). To assess the impact of NOS ES on ICR-associated survival, we conducted a stratified analysis within the NOS ES low and NOS ES high groups. Within the NOS ES low group, there was no significant difference in survival between ICR high and ICR low patients (*P* = 0.66) (top KM curve, **Figure 3F**). However, within the NOS ES high group, the survival difference was highly significant (*P* = 0.001) (bottom KM curve, **Figure 3F)**. These findings were further validated in the TCGA-COAD cohort, where no survival difference was observed in the NOS ES low group (*P* = 0.71) (top KM curve, **Figure 3G**), while a significant difference persisted in the NOS ES high group (*P* = 0.004) (bottom KM curve, **Figure 3G)**. Collectively, these findings suggest that hypermethylation of genes in the NOS pathway suppresses its activity, thereby limiting the survival benefit conferred by immune activation. This points to NOS gene methylation as a potential epigenetic mechanism by which tumors evade effective antitumor immunity.

**Figure 3:**
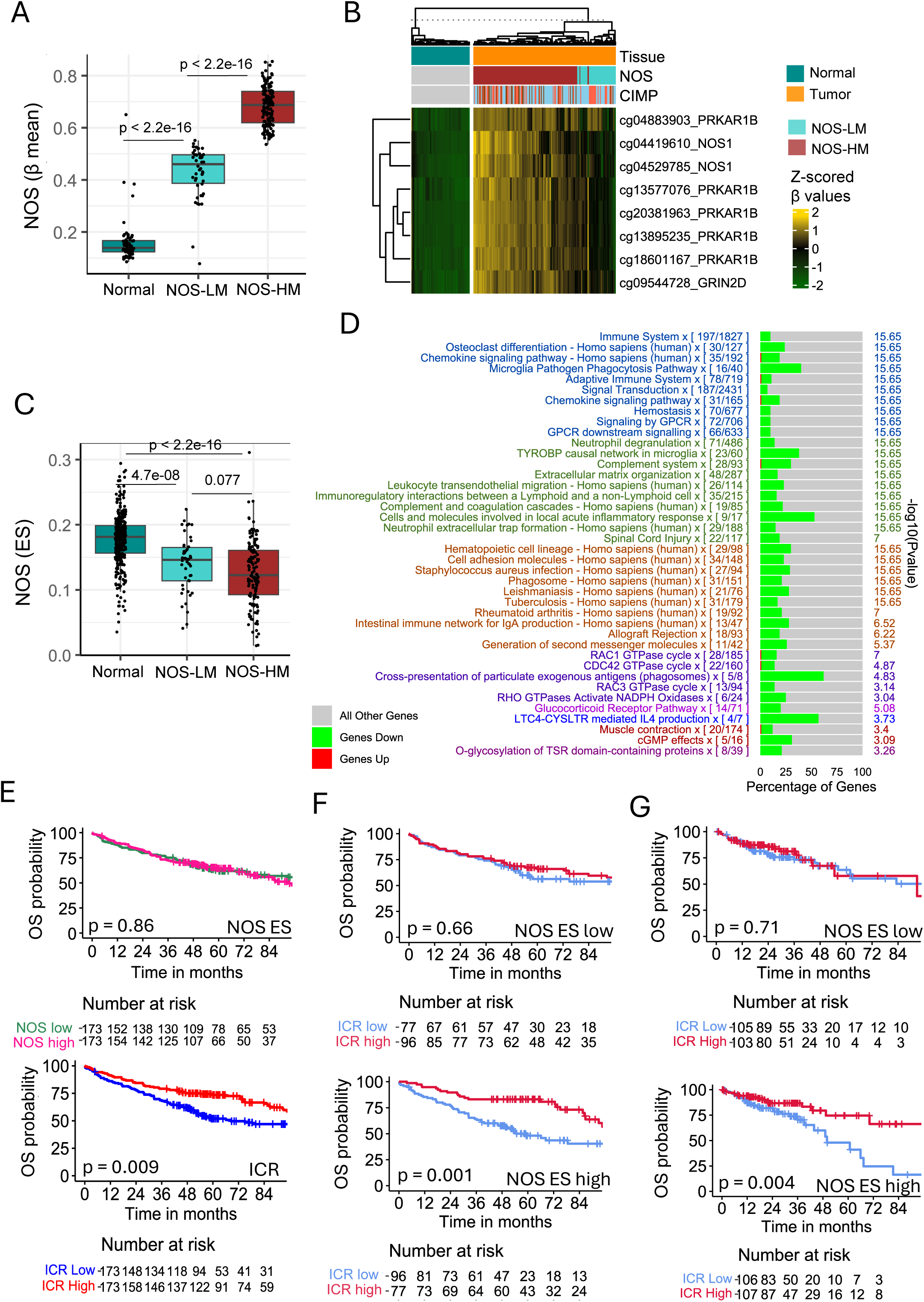
NOS CpG high methylation suppresses immune signaling and modulates ICR-linked survival. **A.** Boxplot showing the β mean of the significant eight DMPs mapped to NOS mapped to genes associated with NOS DMPs from figure 1F for normal, NOS-LM and NOS-HM clusters based on consensus clustering. *P*-value represented from Wilcoxon test. **B.** Heatmap of DMPs (n = 8) mapped to NOS pathway. The heatmap is annotated with tissue, and consensus clusters based on the eight CpGs. **C.** NOS enrichment score (ES) based on genes mapped to the full pathway. *P*-value represented from Wilcoxon test. **D.** Pathway enrichment analysis of DEGs (n = 940, FDR < 0.01, log2FC <= |0.5|) between NOS-HM and NOS-LM. Genes are downregulated/upregulated in NOS-HM vs. NOS-LM. The top 10 pathways from each set are displayed in the bar chart. **E-G.** Kaplan-Meier (KM) plots of overall survival (OS) for: **E.** AC-ICAM cohort patients stratified into high and low groups based on median of NOS ES (top) and ICR score (bottom). **F, G.** ICR groups OS within NOS ES low group (top) and NOS ES high group (bottom) for **F.** AC-ICAM cohort and **G.** TCGA-COAD.

### Biological impact on promoter CpGs

To investigate the overall impact of promoter methylation on gene expression, we analyzed the correlation between promoter CpGs and their mapped genes by performing spearman correlation between gene expression and their promoter’s methylation in tumor samples. Consistent with previous findings (Heery & Schaefer, 2025; Stefansson et al., 2024; VanderKraats et al., 2013), the majority of promoter CpGs (≈ 76%) in our dataset show no significant correlation (p-value > 0.05) with gene expression, reinforcing the understanding that only a subset of promoter methylation events meaningfully impact transcription (pie chart, **Figure 4A**). Furthermore, among CpGs with significant correlation (p-value < 0.05), around 44% have weak spearman coefficient (R = |0.1-0.2|). Among the CpGs with intermediate correlation (R >= |0.5|), 79% are, as expected, negatively correlated with their mapped genes (bar chart, **Figure 4A**).

**Figure 4:**
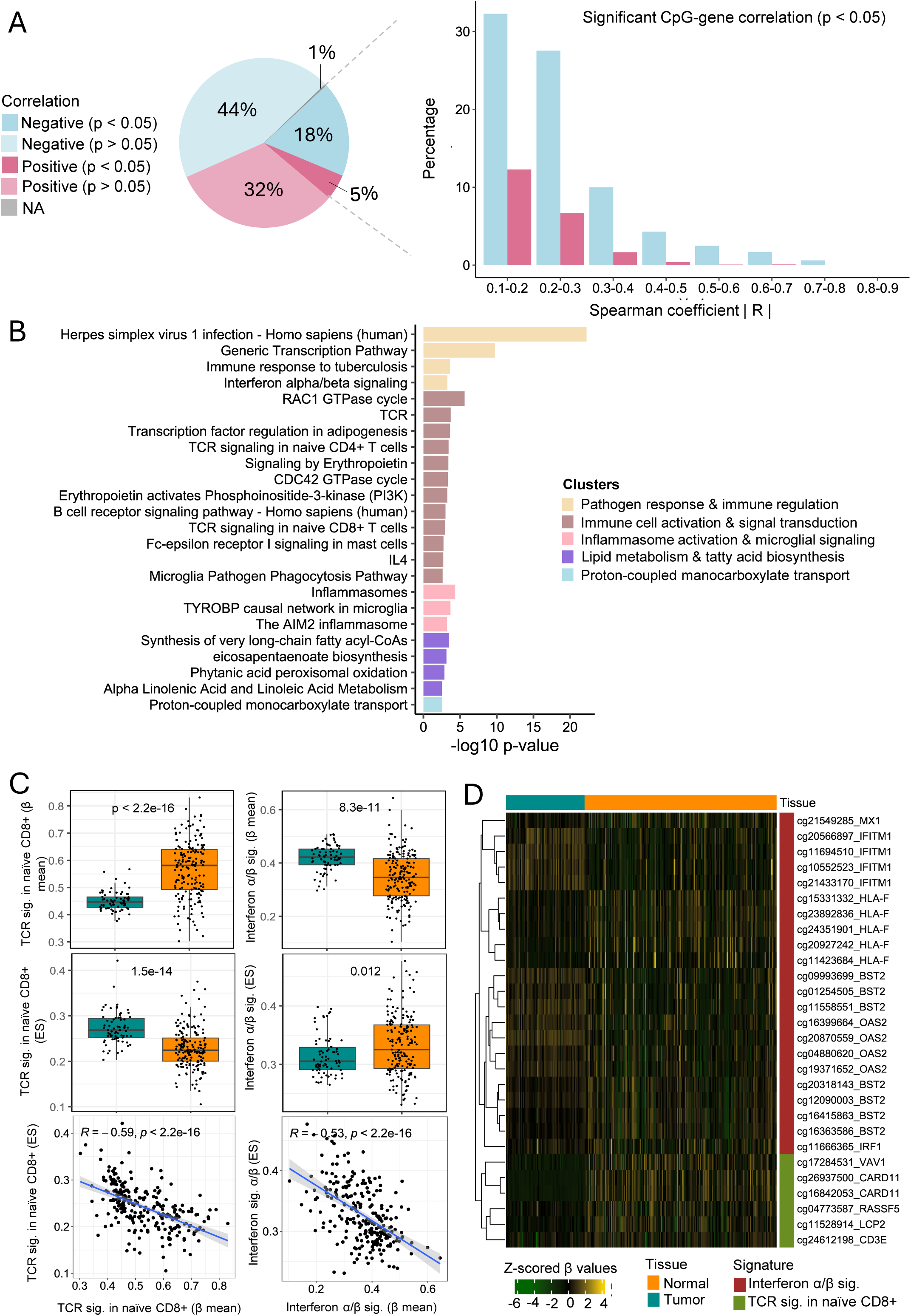
Analysis of promoter CpGs. **A.** Analysis of spearman correlation of promoter CpGs mapped to their genes. Pai chart showing the proportion of significantly negatively correlated CpGs with their mapped genes (18%, *P*-value < 0.05), non-significant negatively correlated CpGs (44%, *P*-value > 0.05), significant positively correlated CpGs (5%, *P*-value < 0.05), non-significant positively correlated CpGs (32%, *P*-value > 0.05). About 1% of the CpGs have no mapped genes, or genes have zero expression data. The barplot shows the correlation of the significant negative and positive correlation (*P*-value < 0.05). **B.** Pathway enrichment analysis of genes (n=260) mapped to negatively correlated promoter CpGs (R <= -0.5, n=899) using CPDB. The plot shows the enrichment of pathways (FDR < 0.05) grouped into five clusters each represented by a different color. The bars indicate the -log *P*-value for each pathway. **C.** Boxplots showing the average β (top plots) of CpGs, and ES (bottom plots) of genes mapped to the pathways identified in B. Scatter plots show the correlation between the ES and average β value. Only results from pathways with significant difference in ES and average β value between tumor and normal and with inverse intermediate correlation (R >= -0.5) are considered. **D.** Heatmap of β value for the CpGs mapped involved in TCR signaling in naïve CD8+ and interferon α/β signaling.

Promoter DNA methylation is a well-established mechanism of transcriptional repression. Thus, to identify CpGs whose methylation status may directly regulate gene expression, we selected CpGs with an intermediate negative correlation for downstream analysis. To gain functional insights, we performed pathway enrichment analysis on the genes (n= 260) mapped to these CpGs (R ≤ -0.5, n = 899). This analysis revealed five distinct clusters associated with key biological processes, including pathogen response and immune regulation, immune cell activation and signal transduction, inflammasome activation and microglial signaling, lipid metabolism and fatty acid biosynthesis, and proton-coupled monocarboxylate transport (**Figure 4B**). To further investigate the impact of epigenetic regulation on these pathways, we calculated an enrichment score (ES) based on gene expression for each pathway included in Figure 4B. Additionally, we computed the mean beta value for CpGs mapped to each pathway to compare between tumor and normal. Moreover, we performed a Spearman correlation analysis between the ES and mean CpG methylation levels of each pathway. We then selected pathways that exhibited an inverse regulatory pattern, where gene expression ES and CpG methylation (mean beta value) showed inverse correlation (R ≤ -0.5). This analysis identified key immune pathways, TCR signaling in naïve CD8+ and interferon α/β signaling that are likely influenced by promoter methylation (**Figure 3C**). For the TCR signaling in naïve CD8+ pathway, promoter CpGs associated with this pathway were significantly hypermethylated in tumors compared to normal colon tissues (higher mean beta value, *P* < 2.2e-15), which corresponded to a significantly lower ES in tumors (*P* = 1.5e-14). The correlation coefficient between mean CpG methylation and ES is -0.59 (*P* < 2.2e-16), suggesting an epigenetic downregulation or reflect a lower immune cell abundance. Conversely, the interferon α/β signaling pathway exhibited hypomethylation in promoter CpGs within tumors (*P* = 8.3e-11), leading to higher ES (*P* = 0.012) compared to normal tissues, with a correlation coefficient of -0.53 (*P* < 2.2e-15). 22 and 6 CpGs were mapped to the interferon α/β signaling and TCR signaling in naïve CD8+, respectively (**Figure D**).

### Interplay between methylation CIMP and microbiome

To investigate whether distinct microbial profiles are associated with different methylation phenotypes, we examined the relationship between methylation status and the relative abundance of bacterial taxa, as previously determined by 16S rRNA gene (141 patients) and WGS (99 patients). As an initial step, we compared bacterial composition between the major methylation subgroups, CIMP-H and CIMP-L, at both the genus and species levels. At the genus level, we identified 14 genera that were significantly different between CIMP-H and CIMP-L (p < 0.05), of which five are more abundant in CIMP-H and nine in CIMP-L. The most significant difference with higher abundance in CIMP-H was observed for *Ruminococcus gnavus group* (*P* = 0.003). Consistent with the literature (Tahara et al., 2014), the relative abundance of Fusobacterium was higher in CIMP-H (*P* = 0.005). On the other hand, *Ruminococcaceae NK4A214 group* was significantly more abundant in CIMP-L (*P* = 0.004) (**Figure 5A**). At the species level, six species were significantly (*P* < 0.05) more abundant in CIMP-H compared to CIMP-L, while three were more abundant in CIMP-L. *Roseburia intestinalis* was the most abundant species in CIMP-H (*P* = 0.003), while *Akkermansia muciniphila* is the most significant in CIMP-L (*P* = 0.027) **(Figure 5B)**. Since CIMP-H is associated with MSI-H and right sided tumor, we performed another analysis (supplementary table 8) by excluding MSI-H, the results retained five species from the original comparison and identified four additional species.

**Figure 5:**
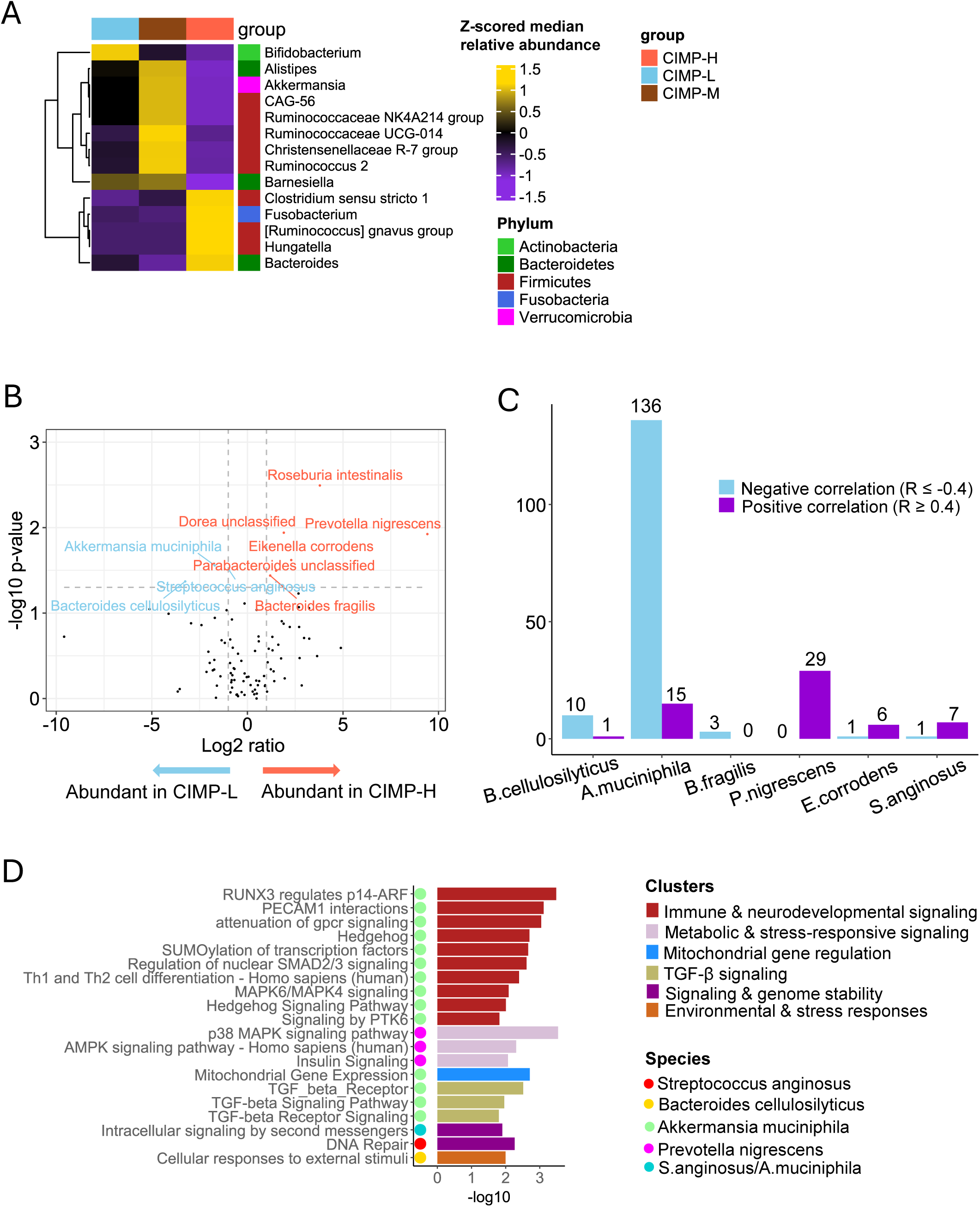
Methylation and microbiome analysis. **A.** Significantly abundant genera between CIMP-H and CIMP-L based on Wilcoxon test (*P*-value < 0.05). Genera that produced NAs after z-scoring were excluded. **B.** Abundant species in CIMP phenotypes. Species with significant difference between CIMP-H and CIMP-L (Wilcoxon *P*-value < 0.05) are highlighted in the volcano plot. **C.** Spearman correlation of promoter CpGs with significantly abundant species from panel B (microbiome unclassified species were excluded). Only CpGs with correlation coefficients R <= | 0.4 | were selected for bar plot illustration. **D.** Pathways enrichment analysis of genes mapped to the selected CpGs for each species from panel C. The bar plot shows enriched pathways (FDR < 0.05) and the species correlated with each cluster. Bars’ colors represent pathways which belong to the same set. Circles represent the species correlated with each pathway.

To better understand how these CIMP-associated bacterial species relate to methylation, we correlated the relative abundance of the identified species (unclassified bacteria were excluded) with promoter CpG methylation levels. This revealed more negative than positive correlations (151 vs 58 CpGs, R =< |0.4|), suggesting that presence of these bacteria is mostly associated with reduced levels of methylation of the identified CpGs (**Figure 5C**). To gain insight into the potentially affected biological pathways, we performed pathway analysis on the genes associated with these CpGs (**Figure 5C**). Pathways enrichment analysis showed six different sets of pathways that were associated with distinct bacterial species. *Akkermansia muciniphila* was associated with two sets of pathways mainly immune and neurodevelopmental signaling and TGF-β signaling. *Prevotella nigrescens* were linked to metabolic & stress-responsive signaling pathways including p38 MAPK and AMPK signaling pathways. The intracellular signaling by second messengers’ pathway from signaling and genome stability cluster was linked to both *Akkermansia muciniphila* and *Streptococcus anginosus*. Interestingly, the latter is also associated with DNA repair pathway (**Figure 5D**). These findings highlight a complex and potentially indirect interplay between distinct bacterial species and host methylation landscapes.

### Development and validation of a methylation-based survival prediction model for colon cancer

To identify methylation markers predictive of overall survival (OS) in colon cancer, we constructed an elastic net–based machine learning model using the AC-ICAM cohort as the training dataset. The model selected 43 promoter CpG sites with both positive and negative regression coefficients associated with survival outcomes (**Figure 6A**). Survival analysis demonstrated a clear separation between the high-risk and low-risk groups defined by the model. In the AC-ICAM training cohort, patients in the high-risk group exhibited significantly worse OS (HR = 7.09, 95% CI: 4.24–11.87, *P* = 9.2×10⁻¹⁴) (top KM, **Figure 6B**), this prognostic trend was successfully validated in the TCGA-COAD cohort (HR = 1.78, 95% CI: 1.06–2.98, *P* = 0.03) (bottom KM, **Figure 6B**). An inverse relationship between the methylation and gene expression has been observed for most CpG–gene pairs such as CpGs mapped to *SLFN13*, *LRRC34*, *NHLRC1*, *INPP5D*, *RNLS*, *MGMT* and *HLA-F* (**Figure 6C**). Gene Ontology (GO) enrichment analysis of genes mapped to the 43 CpGs revealed enrichment in immune-related biological processes, including regulation of immune effector processes, B-cell differentiation, and cytokine-mediated signaling (**Figure 6D**). Enriched GO terms in the molecular function category included Fcγ receptor binding, IL12R activity, and TAP2 binding, while enriched cellular components comprised MHC class II complex and ER-endosome membrane contact sites, indicating strong involvement of antigen presentation and adaptive immune signaling mechanisms (**Figure 6D**). To further explore the biological mechanisms underlying the methylation-derived risk score and its relationship to tumor immunity, we performed a pathway-level correlation analysis integrating the risk score with functional and immune-related signatures. Spearman correlation analysis between the risk score and curated pathways from ConsensusPathDB and immune signatures identified several pathways significantly correlated with the risk score (R ≥ |0.3|). Notably, CREB1 phosphorylation, CD103⁺ T cells, T-cell related signatures (TCR, CD8A), and caspase-mediated apoptosis pathways were negatively correlated with the risk score and showed significant lower enrichment in high-risk group, while PTK6 targets and Josephin domain DUBs showed positive correlation (**Figure 6E**). This indicates that the high-risk group may have reduced T-cell activation, cytotoxicity and immune-mediated apoptotic signaling (tumor cell death). Finally, we assessed the association between the methylation-derived risk score and key molecular and clinical phenotypes. The high-risk group was significantly enriched for MSS tumors (*P* = 0.02), had lower TMB (*P* = 0.01), and exhibited lower ICR (*P* = 5×10⁻⁴). Additionally, the risk score showed a significant increase with advanced tumor stage four (*P* = 0.003), further confirming its clinical relevance (**Figure 6F**). Similar observations were made in TCGA-COAD as well (**Supplementary figure 4A**). The OS difference between methylation risk groups remained significant in stratified analysis within molecular subtypes (MSI, TMB and ICR) in addition to stage (**Supplementary figure 4B**). Furthermore, the methylation risk groups remained significantly associated with improved OS in the Cox multivariate analysis (*P* = 4e-11), together with stage, ICR, MSI and TMB (Supplementary table 9).

**Figure 6:**
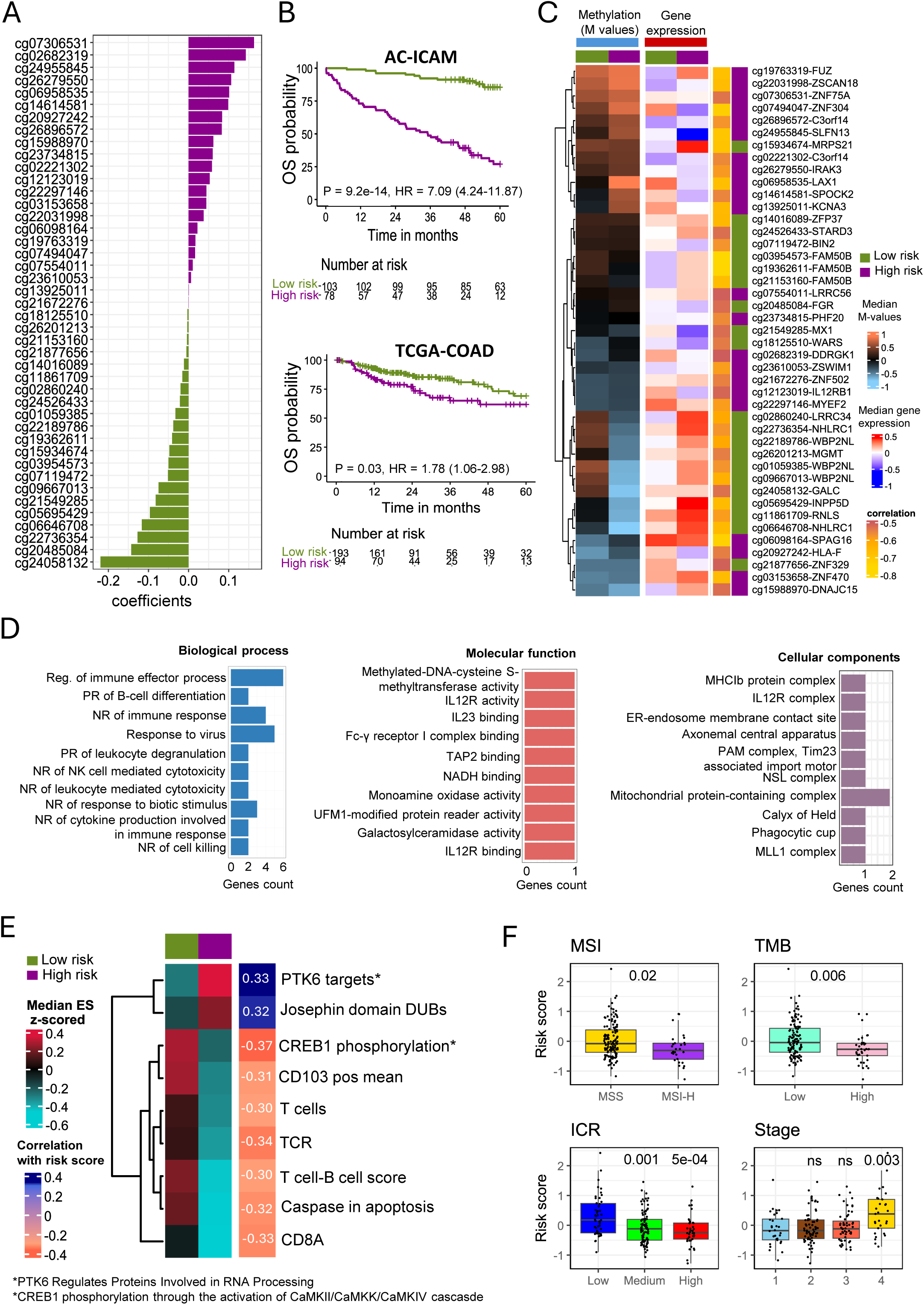
Development and validation of a methylation-based prognostic model for overall survival in colon cancer. **A.** Elastic net regression identified 43 CpG sites significantly associated with overall survival in the AC-ICAM training cohort. Bars represent regression coefficients. **B.** Kaplan–Meier survival curves showing overall five years survival differences between low- and high-risk groups in the AC-ICAM training cohort (top) and TCGA-COAD validation cohort (bottom). Hazard ratios (HRs), 95% confidence intervals, and *P*-values are indicated. **C.** Integrated median heatmaps of methylation (M-values) and gene expression for the 43 CpGs and their mapped genes. Low-risk and high-risk groups are indicated by color bars at the top. Row annotations show the CpG predication (low or high risk) and the correlation between the CpGs and their mapped genes. **D.** Gene Ontology (GO) enrichment analysis of genes associated with the 43 CpGs. Bar plots show enriched terms across Biological Process (BP), Molecular Function (MF), and Cellular Component (CC) categories. Significance cutoff for gene enrichment in a pathway was excluded (FDR = 1, *P*-value =1) to map all genes. **E.** Pathways heatmap showing the ES of selected pathways significantly correlated R ≥ |0.3|) with the risk score. The heatmap displays normalized median ES (z-scored), with row annotations representing the direction and strength of correlation between each pathway and the risk score. **F.** Association of the methylation-based risk score with molecular and clinical phenotypes, including MSI, TMB, ICR, and clinical stage. Boxplots display significant differences in risk scores across groups, with *P*-values from Wilcoxon test indicated (*P* < 0.05 considered significant).

## Discussion

Our previous work on the AC-ICAM cohort provided a comprehensive multi-omics atlas of colon cancer, demonstrating that tumor immune activation, captured by the Immunologic Constant of Rejection (ICR), together with a microbiome signature dominated by *Ruminococcus bromii*, provided strong prognostic information. Building upon this foundation, the current study expands the AC-ICAM dataset by integrating comprehensive methylation profiles from fresh-frozen tumor and matched healthy colon tissues, allowing us to delve deeper into the epigenetic landscape of colon cancer and its interplay with more immune and microbial factors. This integrated approach is crucial for understanding the complex biology of colon cancer, especially given the recognized limitations of existing datasets, such as the TCGA-COAD cohort’s sample inclusion criteria and limited follow-up data.

A central finding of this study is the distinct methylation landscape observed between tumor and normal colon tissues, with tumor samples exhibiting a major shift in their methylation profiles. We identified a substantial number of differentially methylated positions (DMPs), with 63% being hypomethylated and 37% hypermethylated in tumors compared to normal. Notably, hypermethylated DMPs showed a higher proportion of promoter CpGs, confirming previously observed epigenetic gene silencing as a significant mechanism in colon cancer development (Bhootra et al., 2023; M. Li et al., 2023).

Beyond global methylation patterns, we identified specific pathways epigenetically modulated in colon cancer. Extracellular signaling and cell adhesion (ECM-CA) pathways were significantly enriched with DMPs. Intriguingly, ECM-CA-HM tumors exhibited the highest proliferation enrichment score and a downregulation of key immune-related pathways, most notably leukocyte extravasation signaling. This suggests that epigenetic changes affecting the extracellular matrix may contribute to tumor progression by promoting proliferation and suppressing the anti-tumor immune response. This finding is particularly interesting as it proposes another mechanism for how tumors might evade the immune system through epigenetic modifications impacting the tumor microenvironment. While these epigenetic differences clearly influence tumor biology, we observed no statistically significant impact on overall survival between ECM-CA-HM and ECM-CA-LM clusters. This suggests that while ECM-CA methylation changes are biologically relevant, their prognostic utility as a standalone biomarker may be limited, or their effects are modulated by other factors within the complex tumor ecosystem.

Furthermore, our analysis revealed that the nitric oxide signaling (NOS) high methylation cluster (NOS-HM), leads to the downregulation of adaptive immune pathways, including chemokine, TCR, and BCR signaling. Epigenetic silencing of NOS genes may disrupt nitric oxide (NO) production, which is essential for immune cell recruitment and activation. Previous studies have shown that altered NOS signaling can modulate chemokine nitration and reduce T-cell infiltration, thereby promoting an immune-cold tumor microenvironment (Molon et al., 2011; Navasardyan & Bonavida, 2021). Although NOS classification alone did not stratify survival, integrating NOS expression status with the ICR score revealed a strong interaction. In tumors with high NOS expression, ICR-high patients showed significantly better survival, whereas in NOS-low tumors, the prognostic impact of ICR was diminished. This novel finding suggests that NOS plays a role in modulating the immune microenvironment and can influence survival outcomes, particularly by impacting the effectiveness of the anti-tumor immune response captured by ICR. Overall, these findings underscore the importance of considering epigenetic context when assessing the immune landscape of colon cancer.

Investigating the direct impact of promoter methylation on gene expression, we found that while many promoters CpGs showed weak correlations with gene expression, those with intermediate to strong negative correlations supported the concept of transcriptional repression via hypermethylation. Specifically, we identified that TCR signaling in naïve CD8+ related pathways were hypermethylated in tumors, correlating with lower expression of associated genes, potentially indicating an epigenetic mechanism for suppressing T cell activation and subsequent immune rejection. Similar approaches have been used to quantify immune infiltration from methylation data, including a CD8+ TIL signature predictive of prognosis in CRC (Zou et al., 2021). Conversely, the interferon α/β signaling pathway showed promoter hypomethylation and higher expression in tumors, suggesting an active host response or pro-inflammatory TME, though this could also reflect leukocytes infiltration rather than cancer cell-intrinsic changes. These observations provide a deeper understanding of how epigenetic mechanisms might selectively influence immune activity within the tumor microenvironment.

Another aspect of our study explores the interplay between the methylation landscape and the tumor microbiome. We identified distinct bacterial genera and species associated with CIMP-H and CIMP-L phenotypes, for instance, *Ruminococcus gnavus group, Fusobacterium,* and *Roseburia intestinalis* were more abundant in CIMP-H, while *Ruminococcaceae NK4A214 group*, *Akkermansia muciniphila* and *Ruminococcus 2* were more abundant in CIMP-L. Notably, these microbial differences also parallel MSI status and tumor location, consistent with the strong association between CIMP-H and right-sided tumors, suggesting overlapping microbiome signatures across epigenetic and genomic instability subtypes. When we further explored the analysis to account for CIMP-H association with MSI-H and right-sided tumors, five species were retained, suggesting that while part of the observed signal reflects MSI and tumor location, CIMP-associated microbiome differences persist independently. We further reveal that these bacterial species correlate with promoter CpG methylation levels, predominantly showing negative correlations. This is supported by recent work showing that gut microbes can reshape host DNA methylation patterns in colorectal cancer (Z. Liu et al., 2024), and impact tumor suppressor genes such as *Fusobacterium nucleatum* (Xia et al., 2020). Our results expand on this by linking distinct bacterial species with immune-related methylation changes, suggesting that microbiota may directly tune immune surveillance through epigenetic pathways. Remarkably, *Akkermansia muciniphila* was associated with Th1 and Th2 cell differentiation and TGF-β signaling pathway through its correlation with specific CpGs. The role of *A. muciniphila* in colon cancer is increasingly recognized, Fan et al reported its potential beneficial role in cancer immunotherapy and immunomodulatory through enhancing gut barrier integrity, promotes differentiation of regulatory T cells (Tregs), increases expression of anti-inflammatory cytokine IL-10 and TGF-β, and attenuates inflammation in diabetes and colitis models (Fan et al., 2023). Moreover, supplementation with *A. muciniphila* protected mice from CRC progression by specifically inhibiting tryptophan-mediated AhR/β-catenin signaling (Zhang et al., 2023). These findings expand our understanding of the multi-directional influence within the tumor-immune-microbiome axis.

Finally, we developed and validated a robust methylation-based prognostic model that effectively stratifies colon cancer patients into distinct risk groups based on overall survival. The elastic net–derived model, composed of 43 CpG sites, demonstrated a strong predictive performance in both the AC-ICAM training and TCGA-COAD validation cohorts, underscoring its reproducibility across datasets. Integration with gene expression data revealed that many CpG sites correspond to immune-regulatory genes, suggesting that methylation-driven silencing of immune signaling contributes to adverse outcomes in high-risk patients. Functional enrichment analyses indicated that genes linked to the model CpGs are involved in immune effector regulation, antigen presentation, and cytokine receptor signaling pathways essential for tumor immune interactions (Arneth, 2025; Wang et al., 2024). The correlation between the methylation-derived risk score and immune pathway signatures further highlighted that high-risk tumors exhibit diminished T-cell activity, apoptosis signaling, and immune cell infiltration, reflecting an immunologically “cold” phenotype.

Interestingly, among the pathways most positively associated with the high-risk score were PTK6 targets and Josephin domain DUBs, both of which have been implicated in cancer progression. PTK6 (Protein Tyrosine Kinase 6) is a non-receptor tyrosine kinase frequently overexpressed in epithelial cancers, including colorectal carcinoma. PTK6 is highly expressed in colorectal cancer (CRC) tissues and enhances tumor cell proliferation and resistance to chemotherapy in both in vitro and in vivo models. In particular, the activated (phosphorylated) form of PTK6 promotes CRC cell stemness by interacting with and phosphorylating JAK2, thereby triggering the JAK2/STAT3 signaling cascade (C. Liu et al., 2021). Its enrichment in the high-risk group suggests that aberrant PTK6-related signaling contributes to aggressive tumor behavior and poor prognosis. Similarly, deubiquitinating enzymes (DUBs) are proteases responsible for removing ubiquitin moieties from substrate proteins and processing ubiquitin precursors. Although their precise molecular mechanisms remain incompletely defined, numerous DUBs have been implicated in tumor initiation and progression through their regulation of aberrant cancer metabolism, making them promising therapeutic targets in oncology (Tu et al., 2022). Together, these findings suggest that hyperactivation of PTK6 signaling and upregulation of Josephin domain DUBs may serve as key oncogenic mechanisms in methylation-defined high-risk tumors. The concurrent suppression of immune activation and apoptosis pathways in the same group indicates a synergistic effect, whereby oncogenic signaling and immune evasion cooperate to drive poor outcomes. Overall, our results highlight the prognostic and mechanistic value of integrating methylation-based modeling with pathway and immune analyses. The identified CpG signature not only predicts survival but also delineates biologically distinct subtypes, providing a foundation for exploring PTK6 and DUBs as potential therapeutic targets in high-risk colorectal cancers.

In conclusion, this study significantly enriches the AC-ICAM multi-omics atlas by elucidating the intricate epigenetic landscape of colon cancer. We uncover how methylation patterns are interconnected with established molecular subtypes, influence key biological processes such as proliferation and immune suppression, and how specific methylation changes (like those in NOS) can modulate the prognostic impact of immune responses. Moreover, we have identified a compelling interplay between specific bacterial species and the host methylation machinery, suggesting a novel mechanism by which the microbiome might impact tumor biology. The discovery of a robust methylation prognostic signature further strengthens the clinical utility of methylation profiling in colon cancer. Importantly, cfDNA-based methylation biomarkers have recently been described as minimally invasive diagnostic and prognostic tools (Long et al., 2024), suggesting that our tissue-derived CpG panel may have translational potential if validated in liquid biopsy contexts. These findings provide valuable insights into colon cancer biology and highlight new avenues for the discovery of personalized therapeutic approaches and precise prognostic biomarkers, contributing to improved patient management and outcomes.

## Declarations

### Ethics approval and consent to participate

The research was approved by a Dutch and Qatari ethics board of the involved institutions. The research was undertaken to high standards in relation to biosafety and regulatory oversight.

## Data availability

Raw idat files and normalized β matrix for Illumina MethylationEPIC, and BAM files for normal RNA will be publicly available upon acceptance of the manuscript.

## Acknowledgements

This work was supported by the Qatar National Research Fund (JSREP07-010-3-005 awarded to W.H. and NPRP11S-0121-180351 awarded to D.B.) and Sidra Medicine Internal funds (SDR100029, D.B. and W.H.). M.C. was also supported by ‘Associazione Italiana per la Ricerca sul Cancro’ under IG 2018, ID 21846 project awarded to M.C. J.D. was supported by a grant from the Qatar Biomedical Research Institute (VR94). The authors would like to acknowledge the Sidra Medicine research administration team and the Omics Core Facility for providing DNA methylation service using the Illumina Infinium MethylationEPIC BeadChip without whom we could not have performed this work.

## Author’s contribution

E.A, W.H, J.R scientific conceptualization. E.A data analysis. R.M. elastic-net model development. C.R samples preparation for sequencing. E.A, W.H, J.R, R.M wrote and edited the manuscript. H.S data deposition. W.H project supervision and coordination. S.S, R.A, N.E, D.B and all authors reviewed the manuscript.

---

**Supplementary figure 1:**
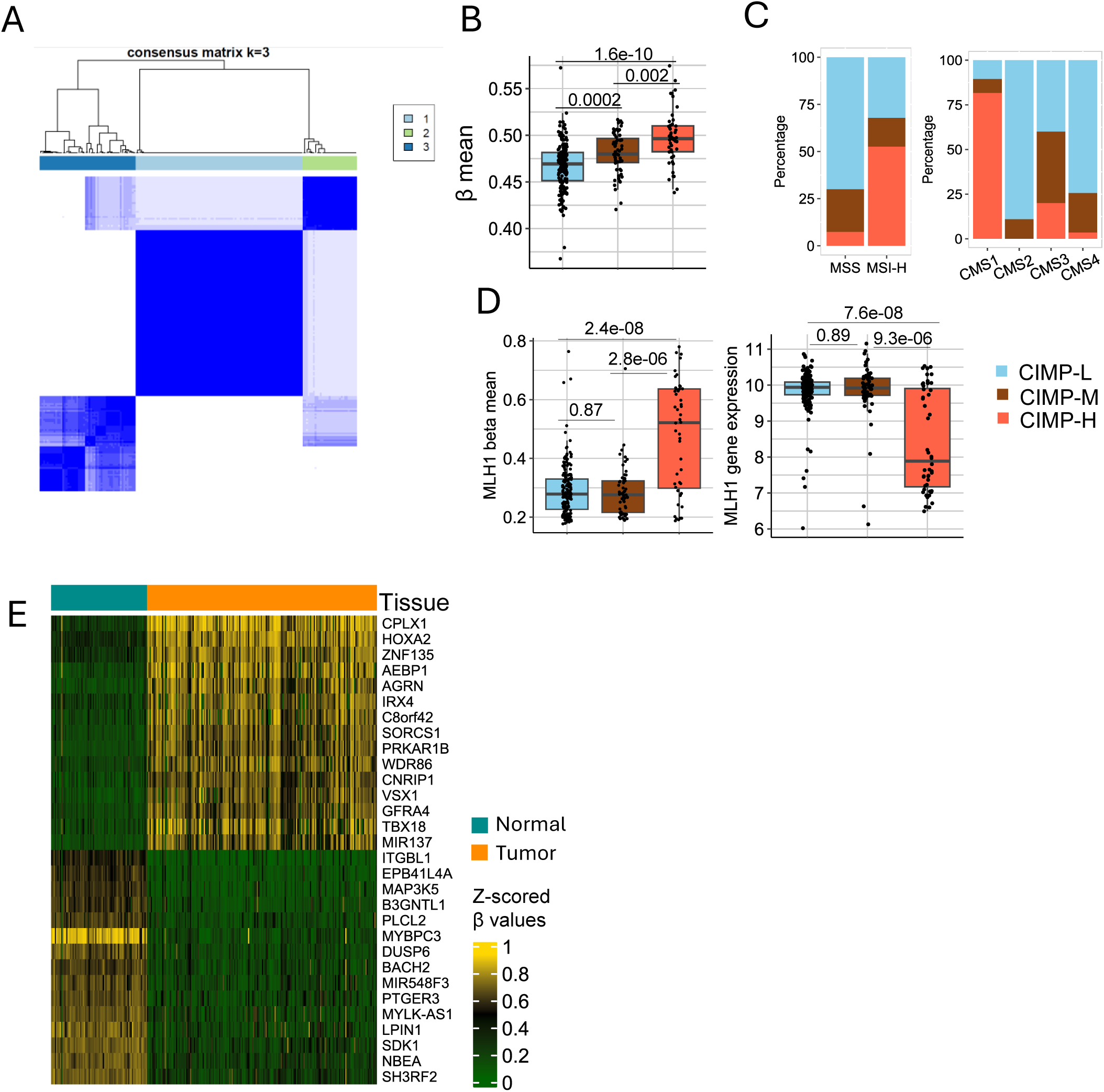
CIMP stratification and molecular associations in TCGA-COAD **A.** Consensus clustering matrix for tumor samples of AC-ICAM to classify samples into different CIMP phenotype based on seven CpGs. **B-D**. CIMP analysis in TCGA-COAD. **B.** Boxplot shows the average β-value of promoter CpGs in CIMP subtypes. CIMP-H has the highest average β-value, while CIMP-L has the lowest. **C.** Stack-bar charts show the frequency of CIMP subtypes in MSI groups (left plot) and CMS subtypes (right plot). **D.** CIMP-H has hypermethylated CpGs of MLH1 genes (left plot), consequently CIMP-H has lower MLH1 gene expression (right plot). **E.** Comparison between hyper- and hypo-methylated genes between tumor and normal in AC-ICAM. The heatmap shows the top 15 hypermethylated and top 15 hypomethylated genes based on the average of all DMPs (FDR < 0.01, Δβ >= |0.4|) mapped to each gene. *P*-value plotted in boxplots from Wilcoxon test.

**Supplementary figure 2:**
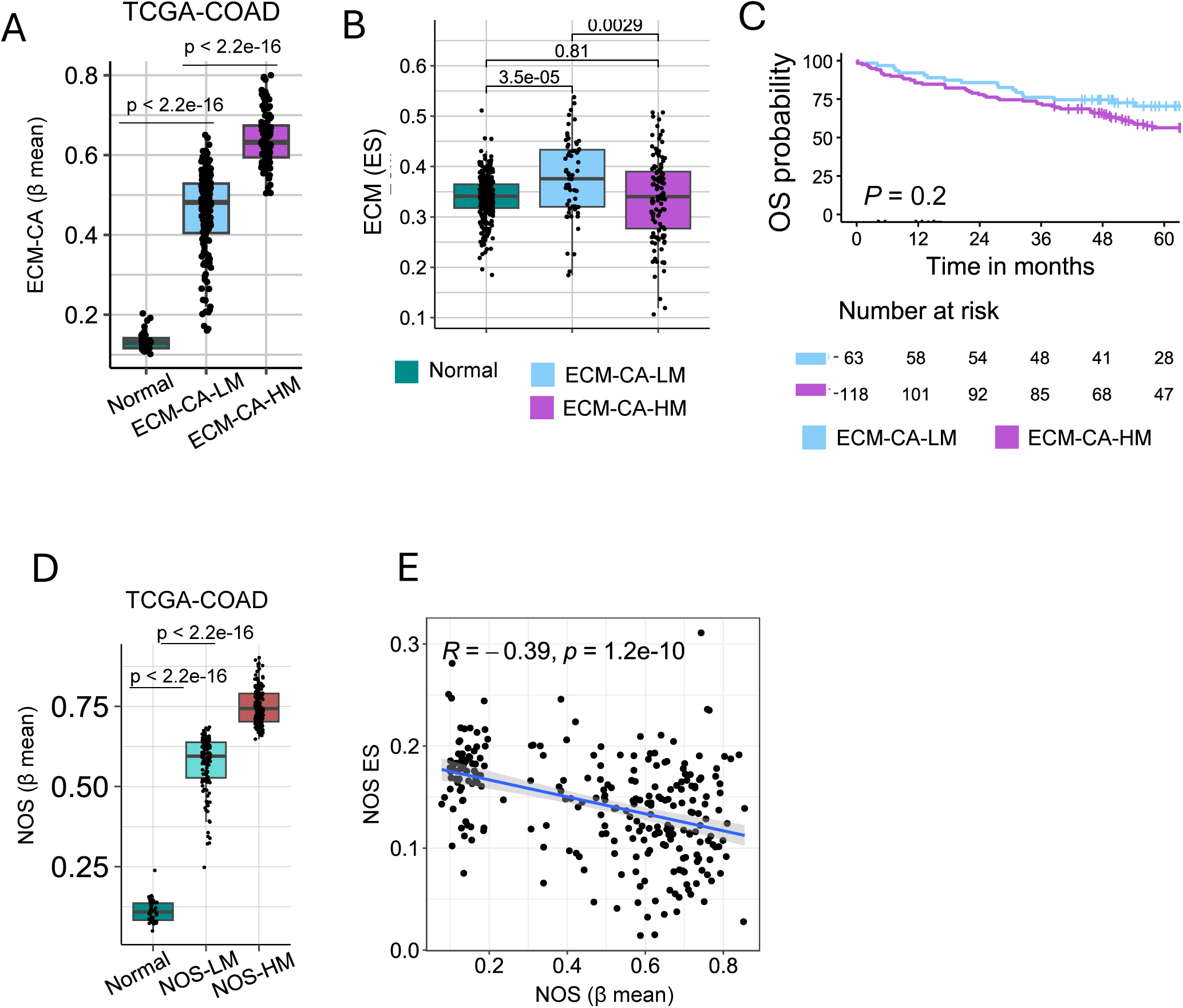
TCGA-COAD validation of ECM-CA and NOS methylation modules **A:** Boxplot shows the 27 CpGs average β-value plotted for normal, ECM-CA-HM and ECM-CA-LM clusters in TCGA-COAD cohort. Plotted *P*-value from Wilcoxon test. **B.** Boxplot shows the extracellular matrix (ECM) ES in normal and ECM-CA clusters. **C.** OS KM plot for ECM-CA clusters in AC-ICAM cohort. **D.** Boxplot showing the β-mean of the significant eight DMPs mapped to NOS in normal and NOS clusters in TCGA-COAD cohort. **E.** Spearman correlation between NOS β-mean and NOS ES in AC-ICAM. Plotted *P*-value in boxplots from Wilcoxon test.

**Supplementary figure 3:**
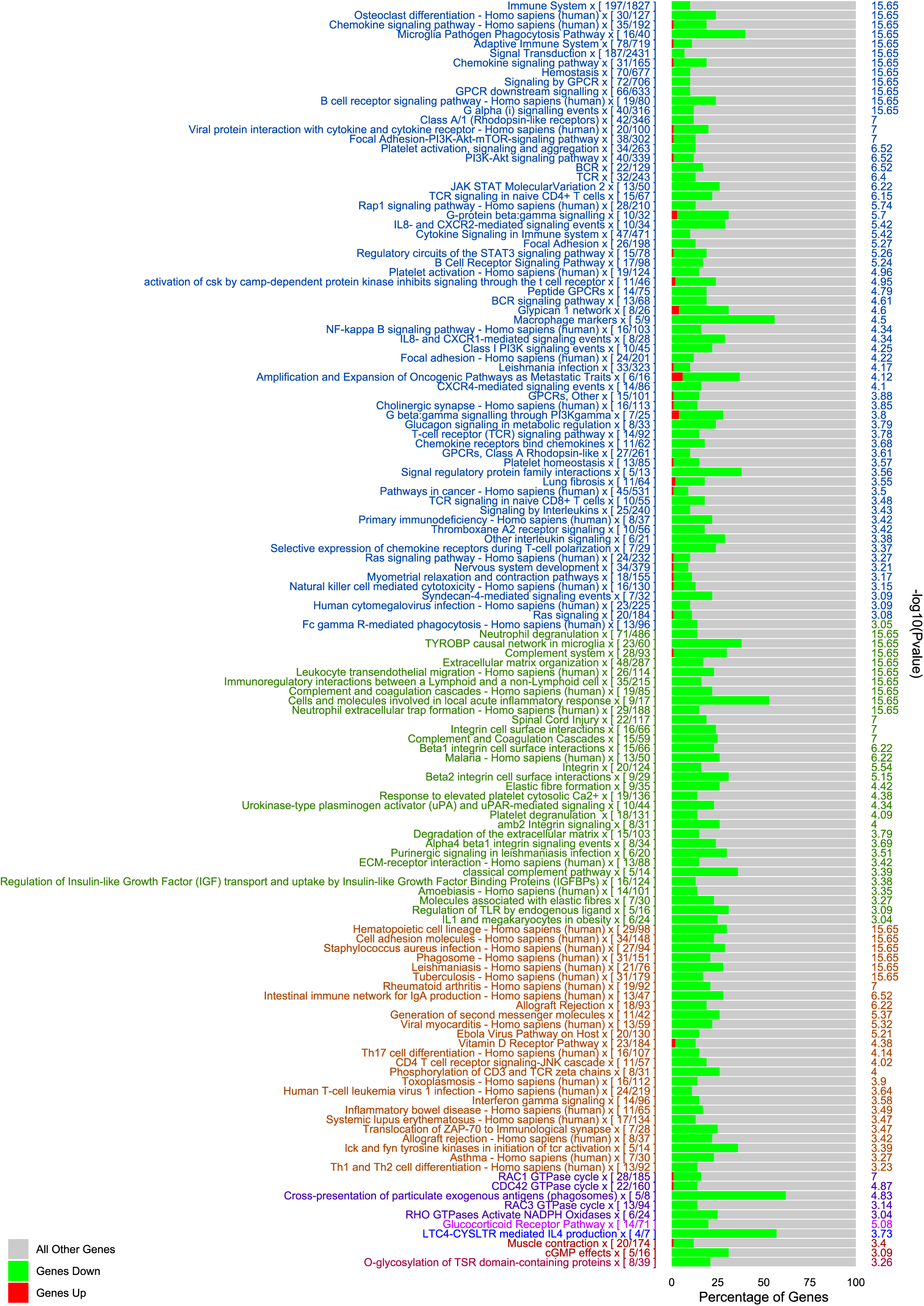
Expanded pathway enrichment for NOS-HM vs NOS-LM, showing broad downregulation of adaptive immune signaling Pathways enrichment analysis of DEGs (n = 940, FDR < 0.01, log2FC <= |0.5|) between NOS-HM and NOS-LM. Genes are downregulated/upregulated in NOS-HM vs. NOS-LM.

**Supplementary figure 4:**
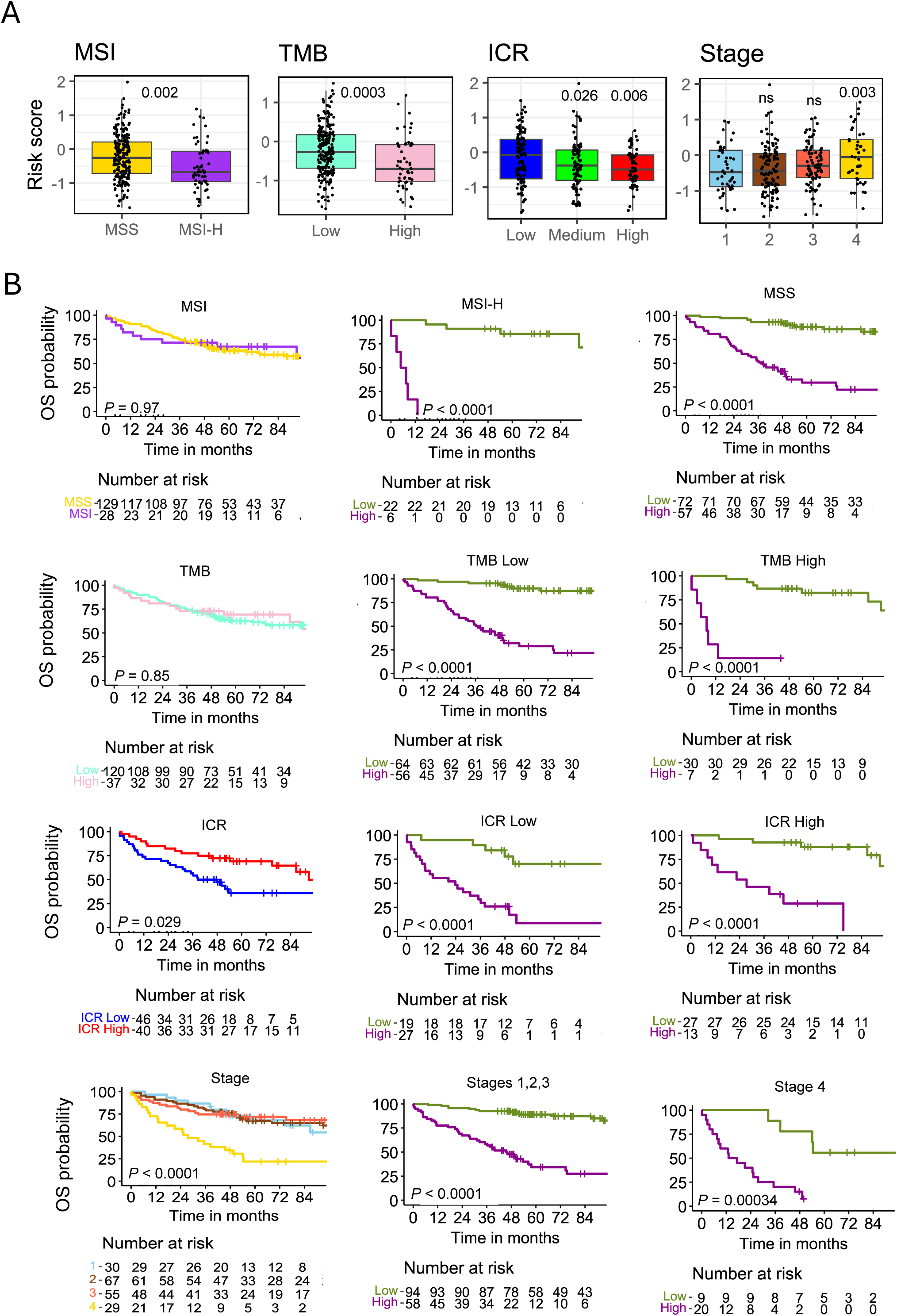
OS ML model validation plots **A.** Association of the methylation-based risk scores with molecular and clinical phenotypes, including MSI, TMB, ICR, and clinical stage in TCGA-COAD cohort. *P*-values from Wilcoxon test are indicated. **B.** OS survival KM plots in AC-ICAM; first column shows the full cohort OS for MSI, TMB, ICR and stage. Second and third columns show stratified OS analysis between low and high-risk groups within those subtypes.

## References

1. Arneth, B. (2025). Molecular Mechanisms of Immune Regulation: A Review. Cells, 14(4), 283. 10.3390/cells14040283

2. Baharudin, R., Ishak, M., Muhamad Yusof, A., Saidin, S., Syafruddin, S. E., Wan Mohamad Nazarie, W. F., Lee, L.-H., & Ab Mutalib, N.-S. (2022). Epigenome-Wide DNA Methylation Profiling in Colorectal Cancer and Normal Adjacent Colon Using Infinium Human Methylation 450K. Diagnostics, 12(1), Article 1. 10.3390/diagnostics12010198

3. Bhootra, S., Jill, N., Shanmugam, G., Rakshit, S., & Sarkar, K. (2023). DNA methylation and cancer: Transcriptional regulation, prognostic, and therapeutic perspective. Medical Oncology, 40(2), 71. 10.1007/s12032-022-01943-1

4. Fan, S., Jiang, Z., Zhang, Z., Xing, J., Wang, D., & Tang, D. (2023). Akkermansia muciniphila: A potential booster to improve the effectiveness of cancer immunotherapy. Journal of Cancer Research and Clinical Oncology, 149(14), 13477–13494. 10.1007/s00432-023-05199-8

5. Gu, S., Lin, S., Ye, D., Qian, S., Jiang, D., Zhang, X., Li, Q., Yang, J., Ying, X., Li, Z., Tang, M., Wang, J., Jin, M., & Chen, K. (2019). Genome-wide methylation profiling identified novel differentially hypermethylated biomarker MPPED2 in colorectal cancer. Clinical Epigenetics, 11(1), 41. 10.1186/s13148-019-0628-y

6. Gutierrez-Angulo, M., Ayala-Madrigal, M., de la L., Moreno-Ortiz, J. M., Peregrina-Sandoval, J., & Garcia-Ayala, F. D. (2023). Microbiota composition and its impact on DNA methylation in colorectal cancer. Frontiers in Genetics, 14. 10.3389/fgene.2023.1037406

7. Heery, R., & Schaefer, M. H. (2025). Systematic identification of regions where DNA methylation is correlated with transcription refines regulatory logic in normal and tumour tissues (p. 2025.02.03.636201). bioRxiv. 10.1101/2025.02.03.636201

8. Janssens, K., Neefs, I., Ibrahim, J., Schepers, A., Pauwels, P., Peeters, M., Van Camp, G., & Op de Beeck, K. (2023). Epigenome-wide methylation analysis of colorectal carcinoma, adenoma and normal tissue reveals novel biomarkers addressing unmet clinical needs. Clinical Epigenetics, 15(1), 111. 10.1186/s13148-023-01516-7

9. Joo, J. E., Mahmood, K., Walker, R., Georgeson, P., Candiloro, I., Clendenning, M., Como, J., Joseland, S., Preston, S., Graversen, L., Wilding, M., Field, M., Lemon, M., Wakeling, J., Marfan, H., Susman, R., Isbister, J., Edwards, E., Bowman, M., … Buchanan, D. D. (2023). Identifying primary and secondary MLH1 epimutation carriers displaying low-level constitutional MLH1 methylation using droplet digital PCR and genome-wide DNA methylation profiling of colorectal cancers. Clinical Epigenetics, 15(1), 95. 10.1186/s13148-023-01511-y

10. Karlsson, S., & Nyström, H. (2022). The extracellular matrix in colorectal cancer and its metastatic settling – Alterations and biological implications. Critical Reviews in Oncology/Hematology, 175, 103712. 10.1016/j.critrevonc.2022.103712

11. Kumar, R. I., Jain, K., Rai, K. R., Gururajan, H., & Sarkar, K. (2025). Targeting epigenetic modifications as an emerging immunotherapeutic strategy for cancers. Immunologic Research, 73(1), 121. 10.1007/s12026-025-09678-7

12. Li, A. H. (2024). Transcriptome Analysis Reveals Anti-Cancer Effects of Isorhapontigenin (ISO) on Highly Invasive Human T24 Bladder Cancer Cells. International Journal of Molecular Sciences, 25(3), 1783. 10.3390/ijms25031783

13. Li, M., Zhu, C., Xue, Y., Miao, C., He, R., Li, W., Zhang, B., Yu, W., Huang, X., Lv, M., Xu, Y., & Huang, Q. (2023). A DNA methylation signature for the prediction of tumour recurrence in stage II colorectal cancer. British Journal of Cancer, 1–9. 10.1038/s41416-023-02155-8

14. Li, Z.-L., Wang, Z.-J., Wei, G.-H., Yang, Y., & Wang, X.-W. (2020). Changes in extracellular matrix in different stages of colorectal cancer and their effects on proliferation of cancer cells. World Journal of Gastrointestinal Oncology, 12(3), 267–275. 10.4251/wjgo.v12.i3.267

15. Liu, C., Pan, Z., Chen, Q., Chen, Z., Liu, W., Wu, L., Jiang, M., Lin, W., Zhang, Y., Lin, W., Zhou, R., & Zhao, L. (2021). Pharmacological targeting PTK6 inhibits the JAK2/STAT3 sustained stemness and reverses chemoresistance of colorectal cancer. Journal of Experimental & Clinical Cancer Research : CR, 40, 297. 10.1186/s13046-021-02059-6

16. Liu, J., Tang, L., Yi, J., Li, G., Lu, Y., Xu, Y., Zhao, S., Mao, R., Li, X., Ren, L., & Wang, K. (2019). Unique characteristics of CpG island methylator phenotype (CIMP) in a Chinese population with colorectal cancer. BMC Gastroenterology, 19(1), 173. 10.1186/s12876-019-1086-x

17. Liu, Z., Zhang, Q., Zhang, H., Yi, Z., Ma, H., Wang, X., Wang, J., Liu, Y., Zheng, Y., Fang, W., Huang, P., & Liu, X. (2024). Colorectal cancer microbiome programs DNA methylation of host cells by affecting methyl donor metabolism. Genome Medicine, 16(1), 77. 10.1186/s13073-024-01344-1

18. Long, Z., Gao, Y., Han, Z., Yuan, H., Yu, Y., Pei, B., Jia, Y., Ye, J., Shi, Y., Zhang, M., Zhao, Y., Wu, D., & Wang, F. (2024). Discovery and Validation of Methylation Signatures in Circulating Cell-Free DNA for the Detection of Colorectal Cancer. Biomolecules, 14(8), 996. 10.3390/biom14080996

19. Molon, B., Ugel, S., Del Pozzo, F., Soldani, C., Zilio, S., Avella, D., De Palma, A., Mauri, P., Monegal, A., Rescigno, M., Savino, B., Colombo, P., Jonjic, N., Pecanic, S., Lazzarato, L., Fruttero, R., Gasco, A., Bronte, V., & Viola, A. (2011). Chemokine nitration prevents intratumoral infiltration of antigen-specific T cells. Journal of Experimental Medicine, 208(10), 1949–1962. 10.1084/jem.20101956

20. Navasardyan, I., & Bonavida, B. (2021). Regulation of T Cells in Cancer by Nitric Oxide. Cells, 10(10), 2655. 10.3390/cells10102655

21. Orecchioni, M., Fusco, L., Mall, R., Bordoni, V., Fuoco, C., Rinchai, D., Guo, S., Sainz, R., Zoccheddu, M., Gurcan, C., Yilmazer, A., Zavan, B., Ménard-Moyon, C., Bianco, A., Hendrickx, W., Bedognetti, D., & Delogu, L. G. (2022). Graphene oxide activates B cells with upregulation of granzyme B expression: Evidence at the single-cell level for its immune-modulatory properties and anticancer activity. Nanoscale, 14(2), 333–349. 10.1039/d1nr04355b

22. Roelands, J., Hendrickx, W., Zoppoli, G., Mall, R., Saad, M., Halliwill, K., Curigliano, G., Rinchai, D., Decock, J., Delogu, L. G., Turan, T., Samayoa, J., Chouchane, L., Ballestrero, A., Wang, E., Finetti, P., Bertucci, F., Miller, L. D., Galon, J., … Bedognetti, D. (2020). Oncogenic states dictate the prognostic and predictive connotations of intratumoral immune response. Journal for Immunotherapy of Cancer, 8(1). 10.1136/jitc-2020-000617

23. Roelands, J., Kuppen, P. J. K., Ahmed, E. I., Mall, R., Masoodi, T., Singh, P., Monaco, G., Raynaud, C., de Miranda, N. F. C. C., Ferraro, L., Carneiro-Lobo, T. C., Syed, N., Rawat, A., Awad, A., Decock, J., Mifsud, W., Miller, L. D., Sherif, S., Mohamed, M. G., … Bedognetti, D. (2023). An integrated tumor, immune and microbiome atlas of colon cancer. Nature Medicine, 1–14. 10.1038/s41591-023-02324-5

24. Stefansson, O. A., Sigurpalsdottir, B. D., Rognvaldsson, S., Halldorsson, G. H., Juliusson, K., Sveinbjornsson, G., Gunnarsson, B., Beyter, D., Jonsson, H., Gudjonsson, S. A., Olafsdottir, T. A., Saevarsdottir, S., Magnusson, M. K., Lund, S. H., Tragante, V., Oddsson, A., Hardarson, M. T., Eggertsson, H. P., Gudmundsson, R.L., … Stefansson, K. (2024). The correlation between CpG methylation and gene expression is driven by sequence variants. Nature Genetics, 56(8), 1624–1631. 10.1038/s41588-024-01851-2

25. Tahara, T., Yamamoto, E., Suzuki, H., Maruyama, R., Chung, W., Garriga, J., Jelinek, J., Yamano, H., Sugai, T., An, B., Shureiqi, I., Toyota, M., Kondo, Y., Estécio, M. R. H., & Issa, J.-P. J. (2014). Fusobacterium in colonic flora and molecular features of colorectal carcinoma. Cancer Research, 74(5), 1311–1318. 10.1158/0008-5472.CAN-13-1865

26. Tu, R., Ma, J., Zhang, P., Kang, Y., Xiong, X., Zhu, J., Li, M., & Zhang, C. (2022). The emerging role of deubiquitylating enzymes as therapeutic targets in cancer metabolism. Cancer Cell International, 22(1), 130. 10.1186/s12935-022-02524-y

27. VanderKraats, N. D., Hiken, J. F., Decker, K. F., & Edwards, J. R. (2013). Discovering high-resolution patterns of differential DNA methylation that correlate with gene expression changes. Nucleic Acids Research, 41(14), 6816–6827. 10.1093/nar/gkt482

28. Wang, C., Chen, L., Fu, D., Liu, W., Puri, A., Kellis, M., & Yang, J. (2024). Antigen presenting cells in cancer immunity and mediation of immune checkpoint blockade. Clinical & Experimental Metastasis, 41(4), 333–349. 10.1007/s10585-023-10257-z

29. Weisenberger, D. J., Siegmund, K. D., Campan, M., Young, J., Long, T. I., Faasse, M. A., Kang, G. H., Widschwendter, M., Weener, D., Buchanan, D., Koh, H., Simms, L., Barker, M., Leggett, B., Levine, J., Kim, M., French, A. J., Thibodeau, S. N., Jass, J., … Laird, P. W. (2006). CpG island methylator phenotype underlies sporadic microsatellite instability and is tightly associated with BRAF mutation in colorectal cancer. Nature Genetics, 38(7), 787–793. 10.1038/ng1834

30. Xia, X., Wu, W. K. K., Wong, S. H., Liu, D., Kwong, T. N. Y., Nakatsu, G., Yan, P. S., Chuang, Y.-M., Chan, M. W.-Y., Coker, O. O., Chen, Z., Yeoh, Y. K., Zhao, L., Wang, X., Cheng, W. Y., Chan, M. T. V., Chan, P. K. S., Sung, J. J. Y., Wang, M. H., & Yu, J. (2020). Bacteria pathogens drive host colonic epithelial cell promoter hypermethylation of tumor suppressor genes in colorectal cancer. Microbiome, 8(1), 108. 10.1186/s40168-020-00847-4

31. Yuan, Z., Li, Y., Zhang, S., Wang, X., Dou, H., Yu, X., Zhang, Z., Yang, S., & Xiao, M. (2023). Extracellular matrix remodeling in tumor progression and immune escape: From mechanisms to treatments. Molecular Cancer, 22(1), 48. 10.1186/s12943-023-01744-8

32. Zhang, L., Ji, Q., Chen, Q., Wei, Z., Liu, S., Zhang, L., Zhang, Y., Li, Z., Liu, H., & Sui, H. (2023). Akkermansia muciniphila inhibits tryptophan metabolism via the AhR/β-catenin signaling pathway to counter the progression of colorectal cancer. International Journal of Biological Sciences, 19(14), 4393–4410. 10.7150/ijbs.85712

33. Zou, Q., Wang, X., Ren, D., Hu, B., Tang, G., Zhang, Y., Huang, M., Pai, R. K., Buchanan, D. D., Win, A. K., Newcomb, P. A., Grady, W. M., Yu, H., & Luo, Y. (2021). DNA methylation-based signature of CD8+ tumor-infiltrating lymphocytes enables evaluation of immune response and prognosis in colorectal cancer. Journal for ImmunoTherapy of Cancer, 9(9). 10.1136/jitc-2021-002671

